# Extracellular DDR1 targeting remodels collagen-rich tumor stroma and local immunity

**DOI:** 10.64898/2026.10.29.762758

**Authors:** Xuan Huang, Xinlin Liu, Litao Huang, Guangjun Jing, Kudelaidi Kuerban, Menghan Cai, Qing Yue, Mei Yu, Shanglin Yang, Yi Zhun Zhu, Li Ye

**Affiliations:** School of Pharmacy and Laboratory of Drug Discovery from Natural Resources and Industrialization, Macau University of Science and Technology, Macau SAR 999078, China; Department of Oncology, Cancer Institute of the Affiliated Hospital of Qingdao University, Qingdao University, Qingdao Cancer Institute, Qingdao 266071, China; Department of Biological Medicines & Shanghai Engineering Research Center of Immunotherapeutics, School of Pharmaceutical Sciences, Fudan University, Shanghai 201100, China; iCareab Biotechnology Co., Ltd., Suzhou 215127, China

## Abstract

Collagen-rich stroma can restrict immune-cell access and sustain immune exclusion in solid tumors, but strategies that directly target extracellular matrix-organizing receptors remain limited. Here, we develop cross-reactive nanobody–Fc proteins against extracellular discoidin domain receptor 1 (DDR1) and identify DDR1nb-3-Fc as a lead molecule with activity in immunocompetent mammary and pancreatic tumor models, including an orthotopic pancreatic model. Tumor control is accompanied by altered collagen architecture and remodeling of the intratumoral immune compartment rather than consistent direct tumor-cell growth inhibition. Single-cell profiling with mouse-level compositional and pseudobulk analyses reveals reproducible pathway-level remodeling of tumor-associated neutrophils, while Ly6G-positive cell depletion significantly modifies the antitumor effect of DDR1nb-3-Fc. These findings identify extracellular DDR1 as an actionable regulator of collagen-rich tumor stroma and support its targeting as a strategy to reshape the tumor microenvironment and promote tumor control.

**Graphical Abstract:** 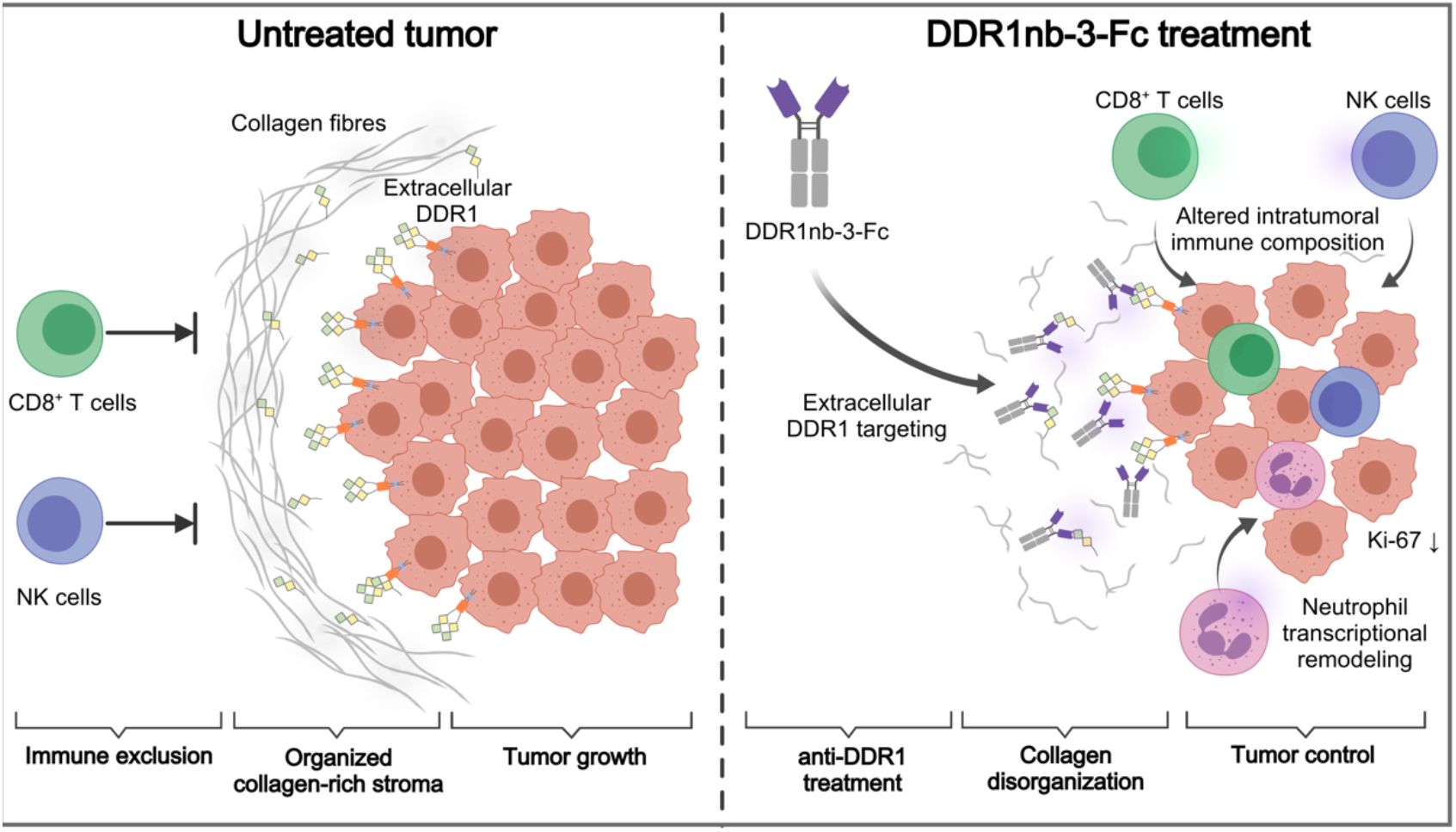

**In brief:** Huang et al. develop cross-reactive nanobody–Fc proteins targeting extracellular DDR1. In immunocompetent mammary and pancreatic tumor models, DDR1 targeting suppresses tumor growth alongside changes in collagen architecture and local immunity. Mouse-level transcriptomic analyses and Ly6G-directed perturbation reveal an additional neutrophil-associated component of the therapeutic response.

**Highlights:** Extracellular DDR1 targeting remodels collagen-rich tumor stroma

DDR1 nanobody–Fc suppresses mammary and pancreatic tumor growth

DDR1 targeting alters intratumoral lymphoid and neutrophil compartments

Ly6G-positive cells modify the response to DDR1nb-3-Fc

## Introduction

The tumor microenvironment (TME) is a major determinant of tumor progression and therapeutic response, and its nonmalignant components increasingly represent actionable targets for cancer treatment (*1-3*). Among these components, the extracellular matrix (ECM) is particularly important because it is not merely a structural scaffold, but an active regulator of tumor behavior and immune-cell trafficking (*2, 4*). In many solid tumors, excessive collagen deposition, crosslinking, and fiber alignment generate a dense and mechanically restrictive stroma that limits immune-cell penetration and contributes to immune exclusion (*4-6*). This feature is prominent in collagen-rich malignancies such as triple-negative breast cancer (TNBC) and pancreatic ductal adenocarcinoma (PDAC), in which stromal barriers can constrain effective antitumor immunity (*7-9*). Therapeutic strategies that remodel collagen architecture may therefore provide a means to improve immune accessibility within otherwise poorly infiltrated tumors.

Discoidin domain receptor 1 (DDR1) is a collagen-activated receptor tyrosine kinase that links tumor cells to the surrounding extracellular matrix and has been implicated in tissue remodeling, fibrosis, and cancer progression (*10-12*). Elevated DDR1 expression has been reported across multiple tumor types and has been associated with aggressive disease and reduced immune infiltration (*11-13*). Although DDR1 contains a canonical intracellular kinase domain, accumulating evidence indicates that its extracellular domain has functions that are highly relevant to stromal immune exclusion. DDR1 extracellular domain (DDR1-ECD) can be released from the tumor-cell surface and associate with collagen fibers in the surrounding matrix, promoting their organization and alignment (*14, 15*). In breast cancer models, genetic or antibody-mediated interference with DDR1 disrupts collagen organization, increases intratumoral T-cell infiltration, and suppresses tumor growth (*15-17*). DDR1 also contributes more broadly to tumor progression by regulating cancer cell interactions with collagen-rich matrices (*18*). These findings position extracellular DDR1 as a mechanistically distinct therapeutic target at the interface between tumor cells, collagen architecture, and antitumor immunity.

Most pharmacological efforts against DDR1, however, have focused on small-molecule inhibitors of the intracellular kinase domain (*10, 18*). Such compounds can interfere with DDR1 signaling but do not directly target the extracellular DDR1-collagen interaction that contributes to matrix organization. This distinction may be important in tumors in which stromal architecture itself limits immune-cell access. Antibody-based targeting of DDR1-ECD offers a complementary strategy by directly engaging the extracellular component of the receptor and, in principle, interfering with its matrix-associated functions (*19, 20*). However, functionally validated antibodies that recognize both human and mouse DDR1 and can therefore be evaluated across immunocompetent preclinical models remain limited. Establishing such cross-reactive reagents is particularly important for defining how extracellular DDR1 targeting affects tumor growth, collagen architecture, and immune composition in vivo.

Stromal remodeling may also influence immune populations beyond lymphocytes. Neutrophils are highly abundant and functionally plastic components of the tumor immune microenvironment, with both tumor-promoting and antitumor activities reported across different contexts (*21-29*). Single-cell studies have further revealed substantial neutrophil heterogeneity, identifying transcriptionally distinct states associated with inflammation, angiogenesis, antigen presentation, immune activation, and treatment response (*30*). Recent work has shown that effective anticancer therapies can be accompanied by neutrophil states associated with enhanced immune activity and improved responses to immune checkpoint blockade (*31, 32*). These observations raise the possibility that altering the stromal environment may influence not only lymphocyte access, but also the composition and functional state of tumor-associated neutrophils. Whether extracellular DDR1 targeting is associated with such changes has not been well defined.

Here, we developed nanobody-Fc molecules that recognize the extracellular domain of both human and mouse DDR1 and used in vivo efficacy-based screening to identify DDR1nb-3-Fc as a lead candidate. We evaluated extracellular DDR1 targeting in immunocompetent mammary and pancreatic tumor models, including an orthotopic PDAC model, and examined treatment-associated changes in collagen architecture and the immune microenvironment. We further used single-cell profiling and Ly6G-directed perturbation to investigate the neutrophil response associated with DDR1 targeting. Together, this study examines extracellular DDR1 as a therapeutic entry point for remodeling collagen-rich tumor stroma and promoting tumor control.

## Results

### Cross-species and in vivo screening prioritizes DDR1 nanobody–Fc candidates

To enable functional evaluation of extracellular DDR1 targeting in immunocompetent mouse models while retaining relevance to human DDR1, we sought VHHs with cross-species reactivity against the DDR1 extracellular domain (DDR1-ECD). Alpacas were immunized with recombinant human and mouse DDR1-ECD, and serum ELISA confirmed robust reactivity against both antigens (Fig. S1A). VHH repertoires recovered from peripheral-blood mononuclear cells were used to construct phage-and yeast-display libraries, which were subjected to sequential antigen selection, clone screening, and sequencing (Fig. 1A and Figs. S1, B to E, and S2). Ten binding-selected VHHs were subsequently expressed as human IgG1 Fc fusions for functional prioritization (Fig. S3A).

**Figure 1.**
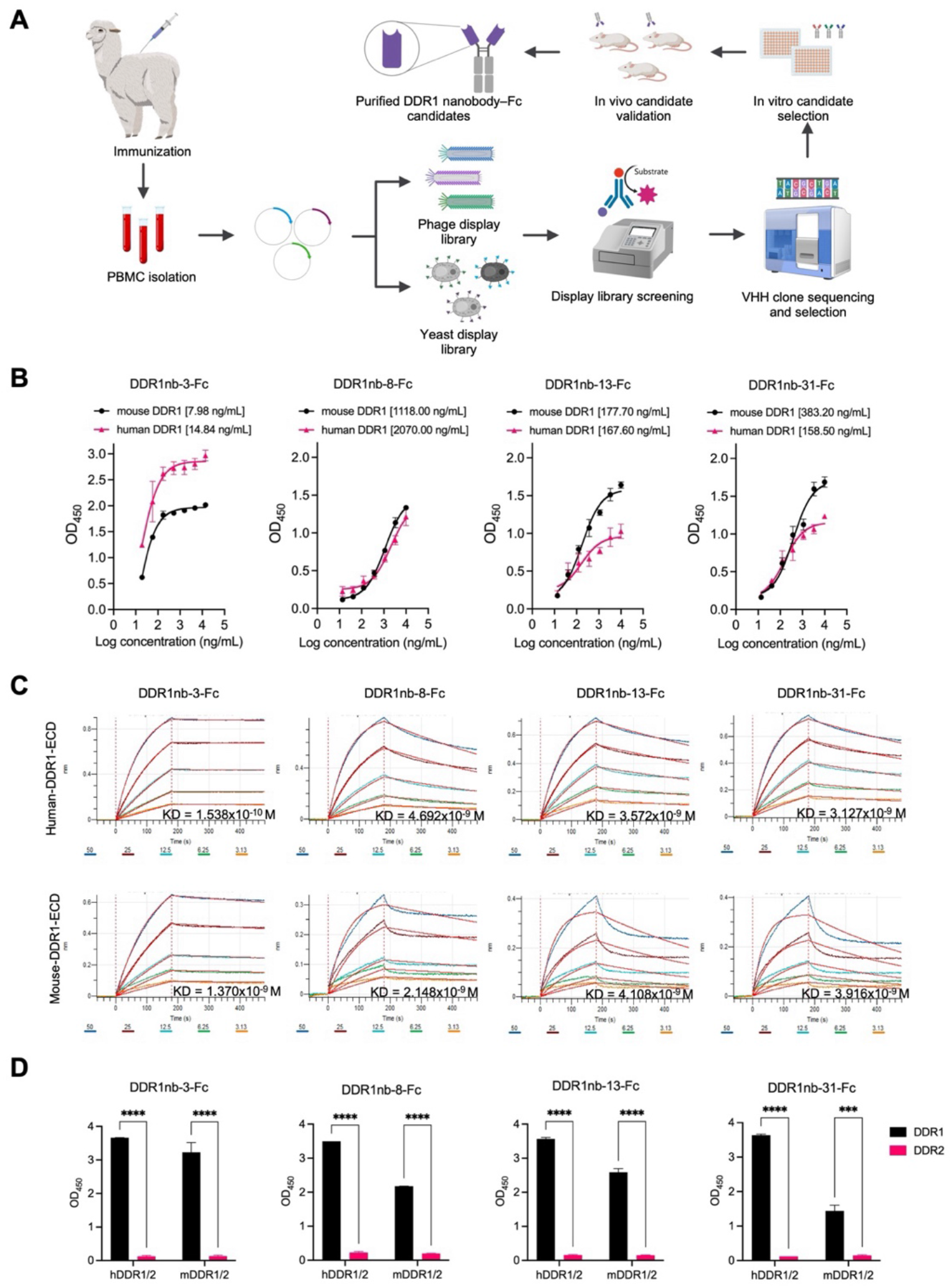
Cross-species binding characterization of four prioritized DDR1 nanobody–Fc candidates. (A) Schematic workflow for alpaca immunization with human and mouse DDR1 extracellular domains (DDR1-ECD), VHH repertoire recovery, phage- and yeast-display selection, sequence screening, nanobody–Fc expression, and candidate prioritization. (B) ELISA binding of DDR1nb-3-Fc, DDR1nb-8-Fc, DDR1nb-13-Fc, and DDR1nb-31-Fc to recombinant human and mouse DDR1-ECD across the indicated concentrations. (C) Representative biolayer interferometry (BLI) sensorgrams showing binding of the four DDR1nb-Fc candidates to human and mouse DDR1-ECD. Apparent kinetic parameters are summarized in Table S1. (D) ELISA comparison of candidate binding to recombinant human or mouse DDR1-ECD and the corresponding DDR2-ECD. OD_450_ values are means ± SD of triplicates. Statistical comparisons were performed using two-way ANOVA. *p < 0.05, **p < 0.01, ***p < 0.001, ****p < 0.0001.

Rather than relying on binding properties alone, we incorporated an early in vivo screen to identify candidates with activity in an immunocompetent tumor setting. In the E0771 subcutaneous model, four of the ten candidates—DDR1nb-3-Fc, DDR1nb-8-Fc, DDR1nb-13-Fc, and DDR1nb-31-Fc—reduced endpoint tumor weight relative to the Fc control (Fig. S3, B to D). These four molecules were therefore prioritized for comparative characterization of cross-species binding, tumor-associated reactivity, and antitumor activity in subsequent studies.

### DDR1nb-3-Fc emerges as a lead candidate with strong cross-species binding and tumor-associated reactivity

We next compared the four prioritized candidates to identify a lead molecule for subsequent mechanistic studies. All four nanobody–Fc proteins bound recombinant human and mouse DDR1-ECD by ELISA and biolayer interferometry (BLI), but DDR1nb-3-Fc consistently showed the strongest cross-species binding profile. It exhibited the lowest ELISA EC50 values, 14.84 ng/ml for human DDR1 and 7.98 ng/ml for mouse DDR1, together with the lowest apparent BLI dissociation constants among the four candidates, 0.1538 nM and 1.370 nM, respectively (Fig. 1, B and C, and Table S1). Binding to the corresponding DDR2 ectodomains was substantially weaker under the same assay conditions (Fig. 1D and Fig. S4), supporting preferential recognition of DDR1 over DDR2.

We then examined whether recombinant-antigen binding translated into reactivity with tumor-associated DDR1. Candidate binding to E0771 cells was strongly influenced by the method used for cell recovery, with higher signals after mechanical scraping than after enzymatic dissociation and the greatest cell-associated reactivity observed for DDR1nb-3-Fc (Fig. 2A).

**Figure 2.**
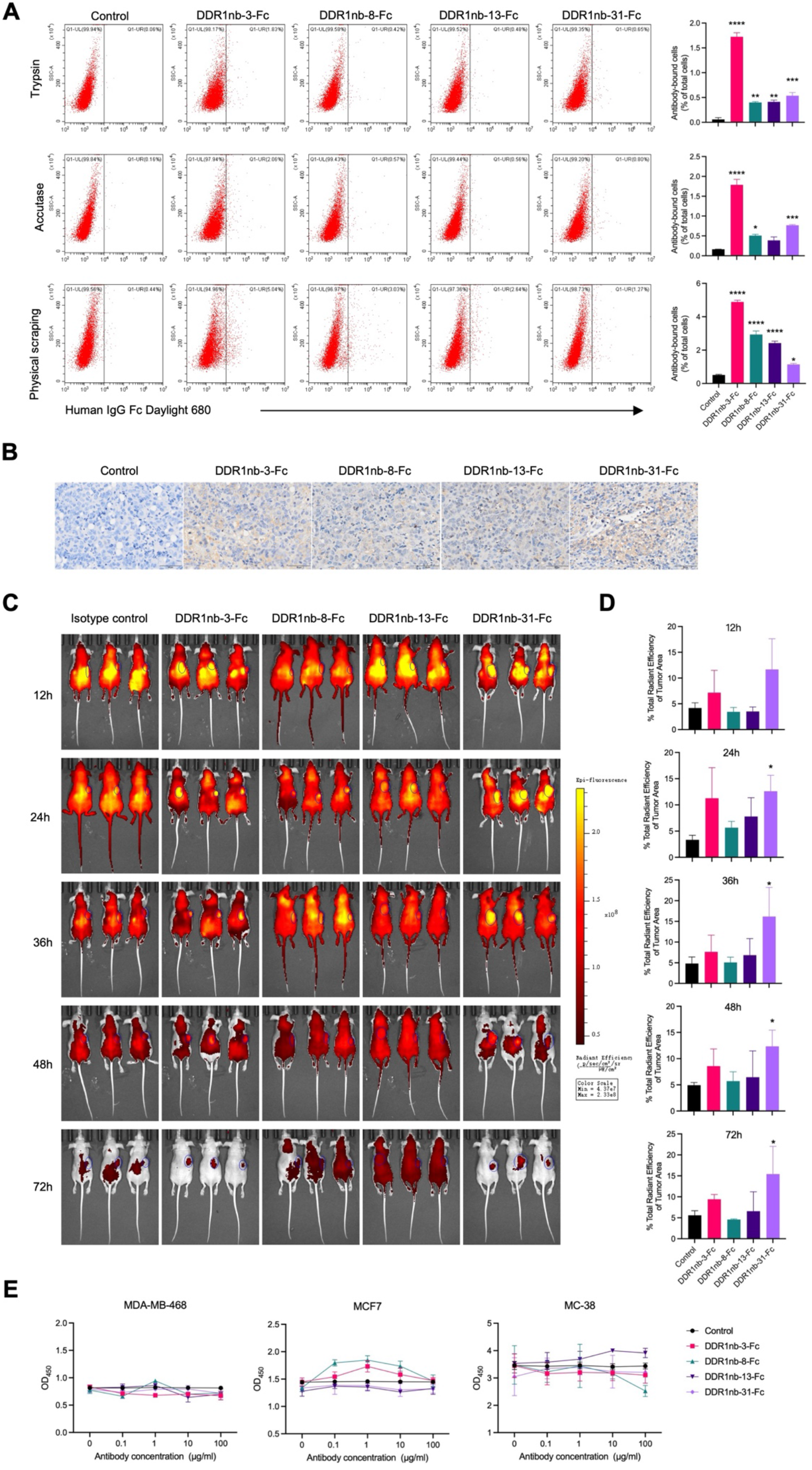
DDR1 nanobody–Fc candidates show tumor-associated reactivity and in vivo localization without direct growth inhibition in vitro. (A) Representative flow-cytometry histograms and quantification of cell-associated binding of the indicated DDR1nb-Fc proteins to E0771 cells collected by trypsinization, Accutase treatment, or mechanical scraping. Quantification is expressed as antibody-bound cells as a percentage of total analyzed cells. Data are presented as mean ± SD from three independent experiments. Statistical comparisons were performed using one-way ANOVA. Asterisks indicate significance versus the control group: *p < 0.05, **p < 0.01, ***p < 0.001, ****p < 0.0001. (B) Representative staining of E0771 tumor sections incubated with the indicated DDR1nb-Fc proteins or human IgG1 Fc control. Scale bar, 50 μm. (C) Representative whole-body fluorescence images of BALB/c nude mice bearing subcutaneous E0771 tumors at 12, 24, 36, 48, and 72 hours after intraperitoneal administration of Cy7-labeled human IgG1 Fc control or the indicated DDR1nb-Fc proteins. (D) Quantification of tumor-associated fluorescence, expressed as the percentage of whole-body fluorescence localized to the tumor region at each time point. Data are presented as mean ± SD (n = 3 mice per group). Statistical comparisons were performed using one-way ANOVA. Asterisks indicate significance versus the control group: *p < 0.05, **p < 0.01, ***p < 0.001, ****p < 0.0001. (E) CCK-8 signal in MDA-MB-468, MCF7, and MC38 cells after 48-hour exposure to the indicated DDR1nb-Fc proteins at the indicated concentrations. Data are presented as mean ± SD (n = 4).

All four candidates also showed stronger staining of E0771 tumor sections than the Fc control (Fig. 2B). We next examined their distribution in E0771-bearing mice using Cy7-labeled proteins. Whole-body imaging revealed visible localization of DDR1nb-3-Fc to the tumor region, with tumor-associated fluorescence remaining detectable at later imaging time points (Fig. 2C). Similar tumor-region signals were observed for the other candidates, with distinct temporal profiles. Quantification of tumor-associated fluorescence relative to whole-body signal further revealed candidate-dependent differences, with DDR1nb-31-Fc showing the most prominent enrichment by this metric, whereas DDR1nb-3-Fc exhibited a more moderate quantitative profile (Fig. 2D). Together, these cellular, tissue, and in vivo imaging data demonstrate that DDR1nb-3-Fc retains tumor-associated reactivity and localization in vivo.

Despite this binding and tumor-associated localization, none of the four candidates produced a consistent concentration-dependent reduction in CCK-8 signal in MDA-MB-468, MCF7, or MC38 cells during 48-hour exposure at concentrations up to 100 μg/ml (Fig. 2E). Thus, the antitumor activity observed during in vivo screening was not reproduced as direct growth inhibition under these culture conditions. Considering its superior cross-species binding profile together with its tumor-associated reactivity and initial in vivo activity, DDR1nb-3-Fc was selected as the principal lead for subsequent studies, while the other three candidates were retained for comparative evaluation.

### DDR1 nanobody–Fc treatment suppresses E0771 tumor growth and is accompanied by collagen remodeling

The absence of consistent growth inhibition in conventional cell culture prompted us to examine whether the in vivo activity of DDR1 nanobody–Fc proteins was associated with changes in the tumor microenvironment. In an independent E0771 efficacy cohort, all four prioritized candidates again slowed tumor growth and reduced endpoint tumor weight relative to the Fc control, confirming the activity observed during discovery-stage screening (Fig. 3, A, C, and D). Body-weight trajectories did not show a progressive treatment-associated decline during the observation period (Fig. 3B).

**Figure 3.**
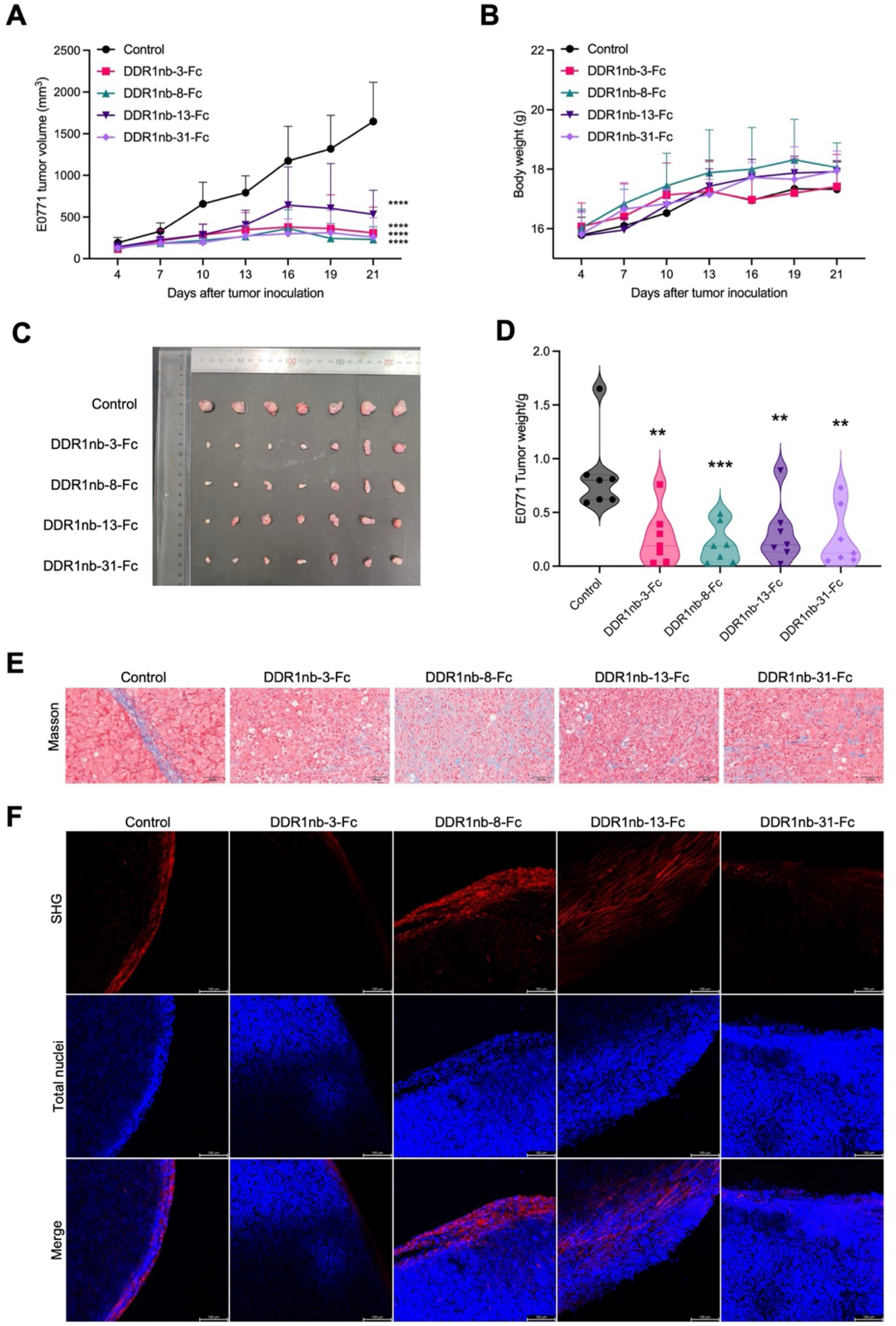
DDR1 nanobody–Fc treatment suppresses E0771 tumor growth and is accompanied by collagen remodeling. (A) Tumor-growth curves in female C57BL/6 mice bearing subcutaneous E0771 tumors and treated with human IgG1 Fc control or the indicated DDR1nb-Fc proteins. Treatment was initiated when tumors reached at least 80 mm³ and administered intraperitoneally at 10 mg/kg twice weekly. (B) Body-weight changes during treatment. (C) Representative images of excised tumors at the study endpoint. (D) Endpoint tumor weights. Points represent individual mice; n = 7 mice per group. (E) Representative Masson’s trichrome-stained sections showing collagen distribution and organization in E0771 tumors from the indicated treatment groups. Scale bars, 50 μm. (F) Representative second-harmonic generation (SHG) images of collagen at tumor margins. SHG collagen signal is shown in red and DCS1 nuclear staining in blue. Scale bars, 100 μm. Data are presented as mean ± SD. Statistical comparisons were performed using one-way ANOVA or two-way ANOVA, as appropriate. Asterisks indicate significance versus the control group: *p < 0.05, **p < 0.01, ***p < 0.001, ****p < 0.0001.

We next examined whether tumor control was accompanied by changes in stromal collagen architecture. Masson’s trichrome staining revealed treatment-associated changes in collagen distribution and organization, characterized by irregular collagen-rich regions rather than a uniform reduction in collagen staining (Fig. 3E and Fig. S5). At the tumor margin, second-harmonic generation imaging showed a continuous collagen-rich boundary in control tumors, whereas this structure appeared less continuous after treatment with DDR1nb-3-Fc or DDR1nb-31-Fc (Fig. 3F). The other candidates produced distinct collagen patterns, indicating that stromal remodeling varied across molecules. Together, these findings link DDR1 nanobody–Fc-mediated tumor control with reorganization of the collagen-rich tumor stroma.

Histopathological examination further showed necrotic and inflammatory regions in treated tumors (Fig. S6). Endpoint serum AST and ALT levels were not increased relative to control, and representative sections of major organs showed no obvious treatment-associated tissue injury (Fig. S7, A to C), providing supportive short-term tolerability observations in this model.

Given the association between collagen-rich stroma and resistance to immune checkpoint therapy, we also evaluated DDR1nb-3-Fc together with PD-1 blockade in a separate E0771 cohort. DDR1nb-3-Fc, anti-PD-1, and the combination each reduced tumor growth relative to control. Factorial analysis of endpoint tumor weights revealed significant main effects of DDR1nb-3-Fc (p = 0.0001) and anti-PD-1 treatment (p < 0.0001), together with a significant DDR1nb-3-Fc × anti-PD-1 interaction (p = 0.0302). The combination produced a lower endpoint tumor weight than DDR1nb-3-Fc alone in the reported comparison (Fig. S8), consistent with an enhanced combined treatment effect. These findings support further investigation of extracellular DDR1 targeting as a stromal-modifying component of PD-1–based immunotherapy.

### DDR1 nanobody–Fc treatment is associated with remodeling of the intratumoral immune compartment

Because collagen-rich stromal architecture can influence immune-cell access, we next asked whether the stromal changes associated with DDR1 targeting were accompanied by alterations in the intratumoral immune compartment. Flow cytometry revealed candidate-specific changes in lymphoid populations. The frequency of CD3-positive cells was increased in tumors treated with DDR1nb-13-Fc or DDR1nb-31-Fc, whereas CD45-, CD4-, and CD8-positive frequencies did not differ significantly from control in the displayed comparisons (Fig. 4A). Tissue staining provided complementary evidence of immune changes within the tumor microenvironment. CD45 and CD8 H-scores were higher after treatment with DDR1nb-3-Fc, DDR1nb-8-Fc, or DDR1nb-31-Fc, and NK1.1 staining was increased in the DDR1nb-3-Fc group (Fig. 4, B and C). In the spleen, CD4- and CD8-positive cell frequencies were increased primarily in the DDR1nb-31-Fc group (Fig. S9), with representative CD4/CD8 flow-cytometry profiles shown in Fig. S10A. Together, these measurements indicate that immune remodeling was not uniform across candidates or compartments, but was most consistently observed within the tumor microenvironment after DDR1nb-3-Fc and DDR1nb-31-Fc treatment.

**Figure 4.**
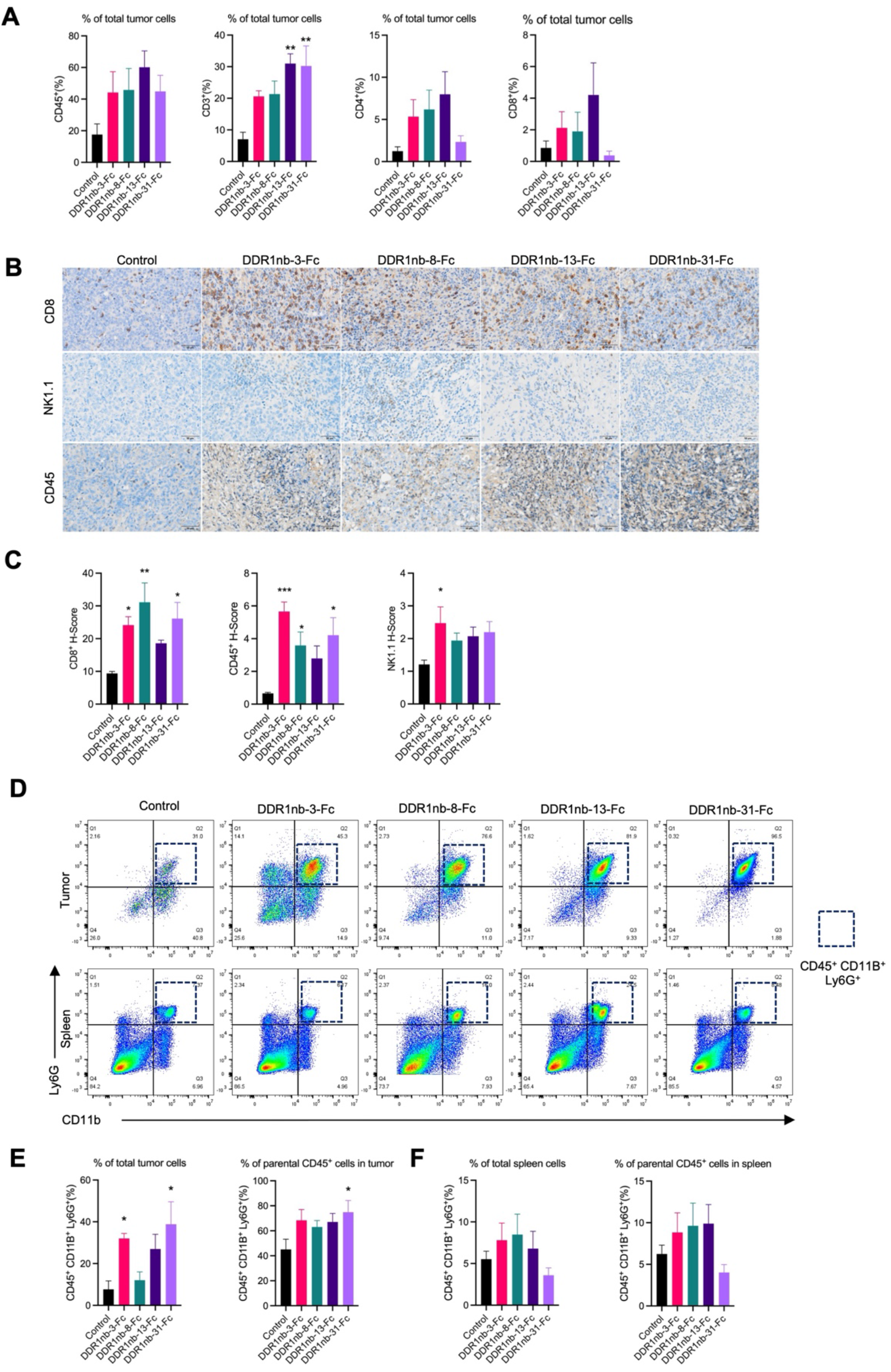
DDR1 nanobody–Fc treatment is associated with candidate-specific changes in intratumoral lymphoid and Ly6G-positive populations. (A) Flow-cytometric quantification of CD45-positive, CD3-positive, CD4-positive, and CD8-positive cells in E0771 tumors. Frequencies are expressed as percentages of total analyzed tumor cells. (B) Representative immunohistochemical staining for CD8, NK1.1, and CD45 in E0771 tumors from the indicated treatment groups. Scale bars, 50 μm. (C) Quantification of CD8, NK1.1, and CD45 immunohistochemical staining using H-scores. (D) Representative flow-cytometry plots showing CD45-positive CD11b-positive Ly6G-positive cells in E0771 tumors and spleens. (E) Quantification of CD45-positive CD11b-positive Ly6G-positive cells in E0771 tumors, expressed either as a percentage of total analyzed tumor cells or as a percentage of parental CD45-positive cells. (F) Quantification of CD45-positive CD11b-positive Ly6G-positive cells in spleens, expressed either as a percentage of total analyzed spleen cells or as a percentage of parental CD45-positive cells. Data are presented as mean ± SD (n = 5–7 mice per group, as indicated). Statistical comparisons were performed using one-way ANOVA. Asterisks indicate significance versus the control group: *p < 0.05, **p < 0.01, ***p < 0.001, ****p < 0.0001.

We also observed changes in the intratumoral Ly6G-positive compartment. The frequency of CD45-positive CD11b-positive Ly6G-positive cells among all analyzed tumor cells was higher in the DDR1nb-3-Fc and DDR1nb-31-Fc groups, whereas within the CD45-positive compartment the increase reached significance only for DDR1nb-31-Fc (Fig. 4, D and E). Corresponding splenic frequencies were not significantly altered (Fig. 4F). Dual Ly6G/CD62L staining further localized neutrophil-associated signals to inflammatory and necrotic regions within treated tumors (Fig. S10B). These findings suggest that DDR1 targeting was associated with local changes in Ly6G-positive cells within the tumor rather than a uniform increase across peripheral immune compartments.

Systemic inflammatory markers also showed a nonuniform response. Endpoint serum IL-6 and IFN-γ concentrations were lower across the four DDR1nb-Fc treatment groups, whereas TNF-α and IL-1β varied among candidates; tumor-associated TNF-α and IL-1β measurements showed distinct patterns from their circulating counterparts (Fig. S7, D to I). Thus, the immune effects associated with DDR1 nanobody–Fc treatment were compartment- and candidate-dependent rather than reflecting generalized systemic immune activation.

### DDR1nb-3-Fc extends tumor control to pancreatic cancer models with associated stromal and immune changes

Having observed tumor control together with stromal and immune changes in the E0771 model, we next asked whether these effects extended to pancreatic cancer. In the subcutaneous KPC1199 model, DDR1nb-3-Fc, DDR1nb-8-Fc, and DDR1nb-31-Fc significantly suppressed tumor growth relative to the Fc control, whereas the DDR1nb-13-Fc comparison did not reach significance (Fig. 5A). Body-weight trajectories showed no progressive treatment-associated decline during the observation period (Fig. 5B). These results demonstrated that the antitumor activity of selected DDR1 nanobody–Fc candidates was not restricted to the mammary tumor setting.

**Figure 5.**
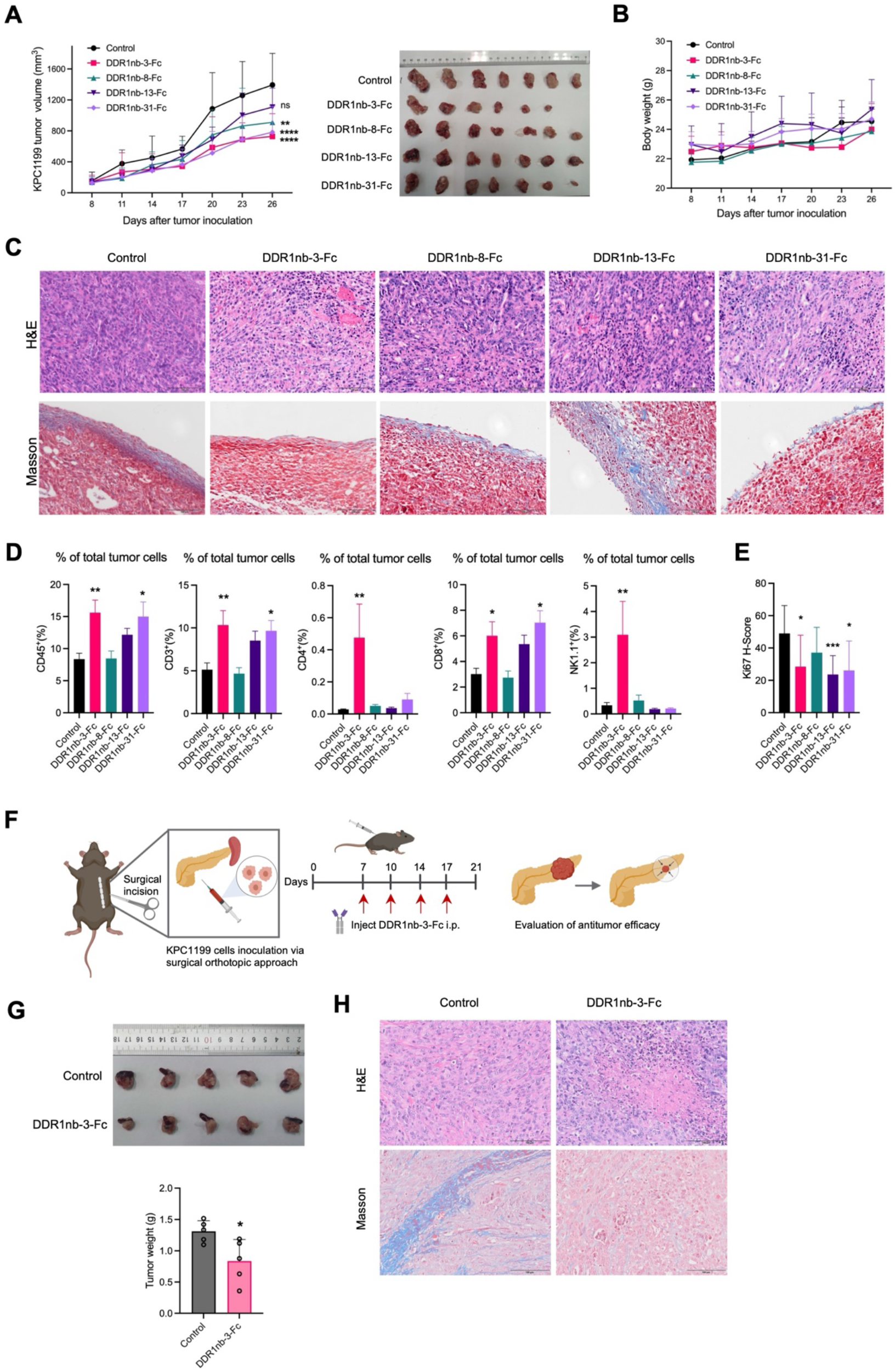
DDR1nb-3-Fc shows antitumor activity with stromal and immune changes in pancreatic cancer models. (A) Tumor-growth curves and representative images of excised tumors from male C57BL/6 mice bearing subcutaneous KPC1199 tumors and treated with human IgG1 Fc control or the indicated DDR1nb-Fc proteins (10 mg/kg, intraperitoneally, twice weekly). n = 7 mice per group. (B) Body-weight changes during treatment in the subcutaneous KPC1199 model. (C) Representative H&E and Masson’s trichrome-stained sections of subcutaneous KPC1199 tumors from the indicated treatment groups. Scale bars, 100 μm. (D) Flow-cytometric quantification of intratumoral CD45-positive, CD3-positive, CD4-positive, CD8-positive, and NK1.1-positive populations. Frequencies are expressed as percentages of total analyzed tumor cells. (E) Quantification of Ki67 immunohistochemical staining using H-scores at the study endpoint. (F) Schematic of the orthotopic KPC1199 implantation and treatment protocol. Six-week-old male C57BL/6 mice received 1 × 10⁶ KPC1199 cells in 50 μl of 1:1 PBS/Matrigel into the pancreatic tail. DDR1nb-3-Fc or human IgG1 Fc control was administered intraperitoneally at 10 mg/kg twice weekly beginning on day 7 after tumor implantation. (G) Representative images of excised orthotopic KPC1199 tumors and quantification of endpoint tumor weights. Points represent individual mice; n = 5 mice per group. (H) Representative H&E and Masson’s trichrome-stained sections of orthotopic KPC1199 tumors from control and DDR1nb-3-Fc-treated mice. Scale bars, 100 μm. Data are presented as mean ± SD. Statistical comparisons were performed using one-way ANOVA or two-way ANOVA, as appropriate. Asterisks indicate significance versus the Fc control where shown: *p < 0.05, **p < 0.01, ***p < 0.001, ****p < 0.0001.

We then examined whether tumor control in KPC1199 tumors was accompanied by changes in the local tumor microenvironment. Representative H&E and Masson’s trichrome staining showed treatment-associated alterations in tumor morphology and regional collagen organization (Fig. 5C). Flow cytometry further revealed higher intratumoral CD45-, CD3-, and CD8-positive cell frequencies in the DDR1nb-3-Fc and DDR1nb-31-Fc groups, while CD4- and NK1.1-positive populations were increased primarily after DDR1nb-3-Fc treatment (Fig. 5D and Fig. S11). Ki67 H-scores were also lower in tumors treated with DDR1nb-3-Fc, DDR1nb-13-Fc, or DDR1nb-31-Fc (Fig. 5E). Thus, as in the E0771 model, tumor control in KPC1199 tumors was accompanied by changes in stromal architecture and intratumoral immune composition, with DDR1nb-3-Fc showing the broadest overall phenotype.

Peripheral immune changes were less uniform. DDR1nb-3-Fc increased splenic CD3- and NK1.1-positive frequencies, whereas DDR1nb-13-Fc reduced several splenic lymphoid populations and DDR1nb-31-Fc reduced the splenic CD8-positive frequency (Fig. S13). Serum TNF-α was increased after DDR1nb-3-Fc or DDR1nb-31-Fc treatment, whereas serum IL-1β did not differ significantly in the displayed comparisons (Fig. S12, B and C). Endpoint organ weights and representative histology provided additional short-term tolerability observations in this model (Fig. S12, A and D to H). These compartment-specific patterns further indicate that the immune effects of DDR1 targeting are not explained by a uniform systemic immune response.

To determine whether DDR1nb-3-Fc retained activity in a more anatomically relevant pancreatic setting, we next evaluated the lead candidate in an orthotopic KPC1199 model, with treatment initiated 7 days after implantation into the pancreatic tail (Fig. 5F). DDR1nb-3-Fc significantly reduced endpoint tumor weight compared with the Fc control (Fig. 5G). Representative H&E and Masson’s trichrome sections showed altered tumor morphology and collagen organization after treatment (Fig. 5H). Endpoint body and liver weights were numerically lower in the DDR1nb-3-Fc group (Fig. S14). Together, these findings extend the antitumor activity of DDR1nb-3-Fc from subcutaneous models to an orthotopic pancreatic tumor setting and further associate extracellular DDR1 targeting with changes in collagen-rich tumor stroma.

### Single-cell and pseudobulk analyses reveal neutrophil transcriptional remodeling, and Ly6G-positive cell depletion modifies DDR1nb-3-Fc-mediated tumor control

To further define the immune landscape associated with DDR1nb-3-Fc treatment in an orthotopic pancreatic tumor setting, we performed single-cell RNA sequencing of CD45-enriched tumor cells from four control and four DDR1nb-3-Fc-treated mice (Fig. 6A). After preprocessing, 57,907 cells were retained and assigned to nine major immune-cell classes (Fig. S15). Neutrophils and mononuclear phagocyte system (MPS) cells represented the largest compartments. At the mouse level, the mean neutrophil frequency was 11.8 percentage points higher in treated tumors, whereas the MPS frequency was 15.3 percentage points lower (Fig. 6B). These differences did not remain significant after correction across the nine immune-cell classes (q = 0.24 and 0.16, respectively), and no other major cell class showed a significant treatment-associated difference (Fig. S15F).

**Figure 6.**
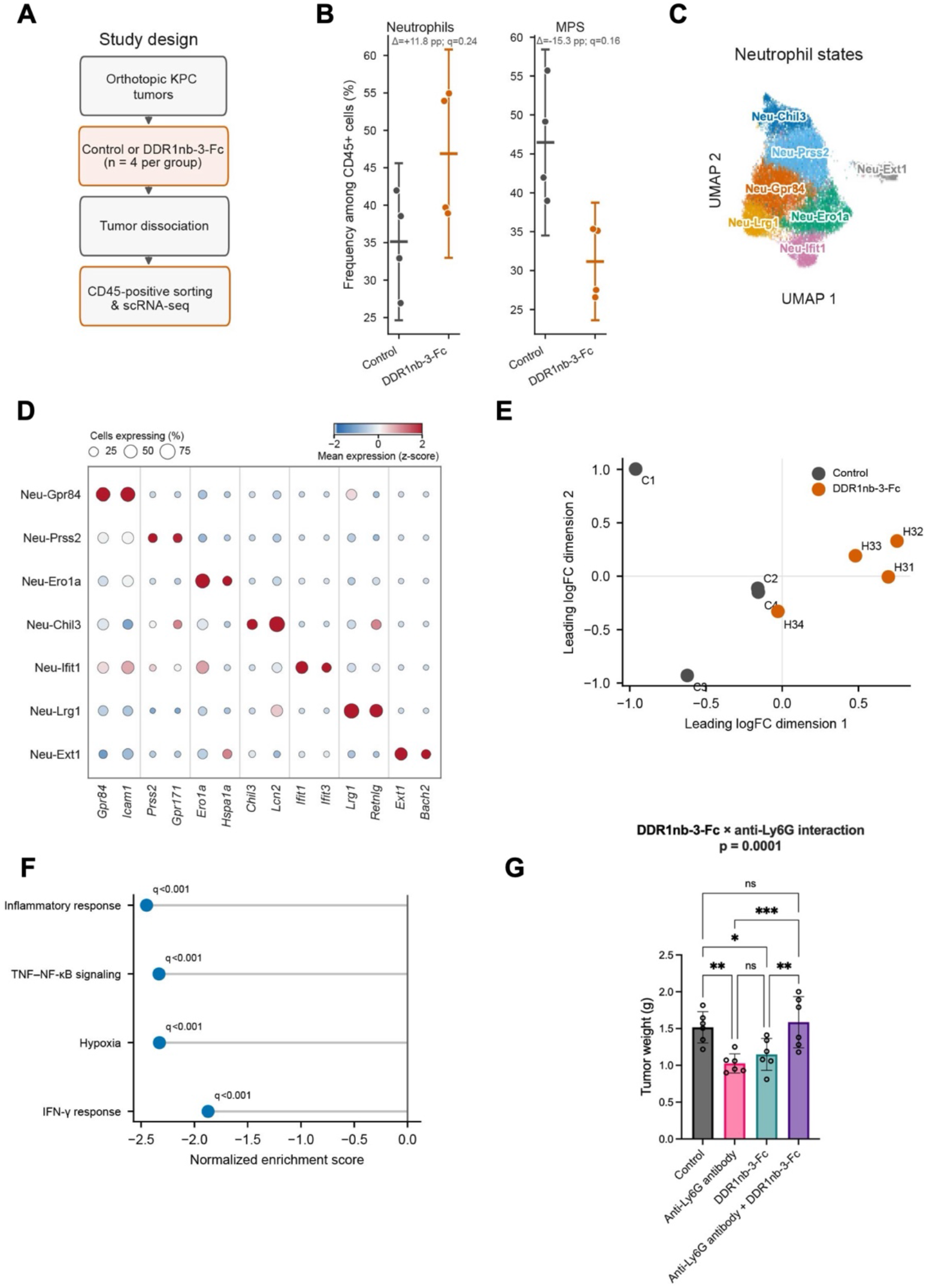
Single-cell and mouse-level pseudobulk analyses reveal treatment-associated neutrophil transcriptional remodeling and evaluate the contribution of Ly6G-positive cells to DDR1nb-3-Fc response. (A) Experimental workflow for single-cell RNA sequencing (scRNA-seq) of CD45-enriched cells isolated from orthotopic KPC1199 tumors following treatment with human IgG1 Fc control or DDR1nb-3-Fc; n = 4 mice per group. (B) Frequencies of neutrophils and mononuclear phagocyte system (MPS) cells among recovered CD45-enriched cells from individual tumors. Points represent individual mice; horizontal lines and whiskers indicate means and 95% confidence intervals (CIs), respectively. (C) UMAP visualization of 22,923 neutrophils assigned to seven transcriptional states. (D) Expression of selected marker genes across the seven neutrophil states. Dot size indicates the percentage of cells expressing each gene, and color indicates z-scored mean log-normalized expression. (E) Multidimensional scaling (MDS) plot of mouse-level whole-neutrophil pseudobulk expression profiles. Each point represents one biological replicate, with control and DDR1nb-3-Fc-treated mice indicated by different colors. (F) Gene-set enrichment analysis (GSEA) of mouse-level whole-neutrophil pseudobulk differential-expression statistics, showing selected Hallmark pathways with significant negative enrichment in DDR1nb-3-Fc-treated tumors relative to controls. Negative normalized enrichment scores (NESs) indicate lower enrichment in the DDR1nb-3-Fc group. Adjusted P values are reported as q values after Benjamini–Hochberg correction. (G) Endpoint tumor weights on day 32 from an independent subcutaneous KPC1199 experiment comparing human IgG1 Fc control, anti-Ly6G antibody, DDR1nb-3-Fc, and DDR1nb-3-Fc plus anti-Ly6G antibody; n = 6 mice per group. For (B), mice were treated as independent experimental units, and cell-class frequencies were compared using two-sided Welch’s t tests with Benjamini–Hochberg correction across the nine major immune-cell classes. For (G), endpoint tumor weights were analyzed using two-way factorial ANOVA with DDR1nb-3-Fc treatment and anti-Ly6G treatment as the two factors. A significant DDR1nb-3-Fc × anti-Ly6G interaction was detected (p = 0.0001). Pairwise comparisons are indicated in the figure. Asterisks denote statistical significance: *p < 0.05, **p < 0.01, ***p < 0.001, ****p < 0.0001.

Given the prominent representation of neutrophils and the Ly6G-positive changes observed in earlier tumor models, we examined the neutrophil compartment in greater detail. Analysis of 22,923 neutrophils resolved seven transcriptional states, designated Neu-Gpr84, Neu-Prss2, Neu-Ero1a, Neu-Chil3, Neu-Ifit1, Neu-Lrg1, and Neu-Ext1 according to their marker profiles (Fig. 6, C and D, and Fig. S16). None of the seven individual state-frequency comparisons reached significance after multiple-testing correction (Fig. S16E). Consistently, global compositional analysis of the seven states did not detect a statistically significant treatment-associated redistribution (exact PERMANOVA p = 0.114, R² = 0.248; Fig. S17A).

We therefore asked whether DDR1nb-3-Fc treatment was associated with transcriptional remodeling of the neutrophil compartment in the absence of a detectable shift in state composition. Raw counts from neutrophils were aggregated at the mouse level to generate eight independent whole-neutrophil pseudobulk samples. Multidimensional scaling was used to visualize mouse-level pseudobulk expression profiles (Fig. 6E). No individual gene reached FDR < 0.05 in the mouse-level differential-expression analysis (Fig. S17B). In contrast, ranked gene-set enrichment analysis identified coordinated negative enrichment of inflammatory response (NES = −2.45, q = 0.0005), TNF–NF-κB signaling (NES = −2.33, q = 0.0005), hypoxia (NES = −2.33, q = 0.0005), and IFN-γ response (NES = −1.87, q = 0.0005) in DDR1nb-3-Fc-treated tumors (Fig. 6F). These pathway-level effects remained directionally consistent across all eight leave-one-mouse-out analyses (Fig. S17C). State-resolved analyses further showed that the transcriptional changes were distributed across multiple neutrophil states rather than being attributable to a single state (Fig. S17D). Together, these analyses support reproducible pathway-level remodeling of the neutrophil compartment without evidence for a dominant treatment-associated redistribution of a specific neutrophil state.

To test whether Ly6G-positive cells were functionally relevant to DDR1nb-3-Fc-mediated tumor control, we performed an independent four-group experiment in the subcutaneous KPC1199 model. Factorial analysis of endpoint tumor weights revealed a significant DDR1nb-3-Fc × anti-Ly6G interaction (p = 0.0001), indicating that Ly6G-positive cell depletion significantly modified the effect of DDR1nb-3-Fc on tumor burden (Fig. 6G). DDR1nb-3-Fc and anti-Ly6G monotherapies each reduced endpoint tumor weight relative to control, whereas co-treatment resulted in greater tumor burden than either monotherapy. Longitudinal tumor-volume measurements showed a corresponding treatment-dependent pattern (Fig. S18A). Consistent with effective Ly6G-positive cell depletion, circulating neutrophil counts were reduced in anti-Ly6G-treated groups at the experimental endpoint (Fig. S18B), whereas organ weights from the same experiment are shown in Fig. S18C.

Together, these findings identify the neutrophil compartment as a transcriptionally responsive component of the DDR1nb-3-Fc-treated tumor microenvironment and provide functional evidence that the Ly6G-positive compartment modifies the therapeutic response to DDR1nb-3-Fc. The absence of a significant global state-composition shift, together with the distributed pathway-level changes across multiple neutrophil states, argues against attributing this effect to a single transcriptionally defined neutrophil population.

## Discussion

Our study identifies extracellular DDR1 as an actionable regulator of the collagen-rich tumor microenvironment. By combining cross-species nanobody screening with functional evaluation in immunocompetent tumor models, we identified DDR1nb-3-Fc as a lead extracellular DDR1-targeting molecule with activity in both mammary and pancreatic tumors. Across these models, tumor control was accompanied by altered collagen architecture and changes in the local immune compartment, whereas consistent direct inhibition of tumor-cell growth was not observed under conventional culture conditions. These findings are consistent with a contribution of the tumor microenvironment to the antitumor activity of extracellular DDR1 targeting, although tumor-cell-autonomous effects in the in vivo models cannot be excluded.

This mode of action distinguishes extracellular DDR1 targeting from conventional approaches directed at DDR1 kinase activity. Most small-molecule DDR1 inhibitors act through the intracellular ATP-binding pocket and primarily interfere with kinase-dependent signaling (*10, 18*). In contrast, extracellular DDR1 can associate with collagen and contribute to collagen-fiber organization independently of canonical kinase signaling (*14, 15*). Previous genetic and antibody-based studies have implicated DDR1 in collagen alignment, immune exclusion, and tumor progression (*17, 33*). Our findings extend this framework using cross-reactive nanobody–Fc proteins that can be evaluated in immunocompetent mouse models while retaining recognition of the human target. The absence of reproducible growth inhibition in vitro, despite preserved DDR1 binding and tumor-associated localization, further suggests that the antitumor activity of these molecules depends substantially on tissue-contextual functions of DDR1. However, because DDR1nb-3-Fc also binds cell-associated DDR1, its activity may involve both matrix-associated and membrane-associated receptor pools.

The stromal effects observed across mammary and pancreatic tumor models further link extracellular DDR1 to matrix-mediated immune regulation. Rather than causing uniform collagen depletion, DDR1 nanobody–Fc treatment was associated with redistribution and altered organization of collagen-rich regions. In E0771 tumors, SHG imaging additionally revealed a less continuous collagen-rich structure at the tumor margin after treatment with selected candidates. Similar treatment-associated changes were observed in subcutaneous and orthotopic KPC1199 tumors. These observations are consistent with previous evidence that DDR1 contributes to collagen-fiber alignment and that antibody-mediated DDR1 targeting can alleviate matrix-associated immune exclusion (*15, 20*). Notably, the four prioritized nanobody–Fc candidates were not equivalent in either stromal or antitumor phenotypes. This divergence suggests that binding affinity alone does not fully determine functional activity and that epitope engagement, receptor context, tissue localization, or other molecular properties may contribute to the biological effects of extracellular DDR1 targeting.

Stromal remodeling was accompanied by changes in the local immune environment. In E0771 tumors, several DDR1 nanobody–Fc treatments increased tissue-associated CD45 and CD8 signals, with additional NK1.1 changes observed after DDR1nb-3-Fc treatment. In KPC1199 tumors, DDR1nb-3-Fc was associated with broader increases in several intratumoral lymphoid populations. By contrast, splenic lymphoid populations and circulating inflammatory mediators showed more heterogeneous responses. This compartmental distinction argues against a uniform systemic immune activation and instead supports a predominantly tumor-localized immune response associated with stromal remodeling. The exploratory PD-1 combination experiment is consistent with this interpretation: DDR1nb-3-Fc and PD-1 blockade showed a significant interaction in the factorial analysis, and the combination reduced endpoint tumor burden relative to DDR1nb-3-Fc alone. This experiment was not designed to establish formal pharmacological synergy, but the result is compatible with the broader concept that reducing matrix-associated constraints can improve the conditions under which antitumor immunity operates (*19*).

The neutrophil compartment emerged as an additional component of the response to extracellular DDR1 targeting. Single-cell profiling resolved seven transcriptionally defined neutrophil states, but neither individual state-frequency comparisons nor global compositional analysis detected a significant treatment-associated redistribution. Instead, mouse-level pseudobulk analysis identified coordinated pathway-level changes involving inflammatory response, TNF–NF-κB signaling, hypoxia, and IFN-γ response programs. These effects were preserved in leave-one-mouse-out analyses and distributed across multiple neutrophil states, indicating that the transcriptional response was not dominated by a single state. Functional perturbation further showed that broad Ly6G-positive cell depletion significantly modified the effect of DDR1nb-3-Fc on tumor burden. Because anti-Ly6G treatment alone also reduced tumor burden, these findings do not support a simple uniformly antitumor or protumor role for Ly6G-positive cells. Rather, they indicate that this compartment contributes to the treatment response in a context-dependent manner during DDR1-directed stromal remodeling.

From a therapeutic perspective, DDR1nb-3-Fc combines several features that facilitate further investigation of extracellular DDR1 biology. Its cross-reactivity with human and mouse DDR1 enables evaluation in immunocompetent preclinical systems while retaining relevance to the human target, and its activity across mammary and pancreatic tumor models suggests that extracellular DDR1 targeting may be applicable beyond a single tumor lineage. More broadly, our findings support a view of collagen-rich stroma as an active regulator of immune accessibility and identify extracellular DDR1 as a point of intervention at the interface between tumor cells, extracellular matrix organization, and local immunity. These results provide a rationale for further development of extracellular DDR1-directed biologics and for testing their integration with therapies that depend on effective immune-cell access to tumors.

## Limitations of this study

Several aspects of extracellular DDR1 targeting remain to be resolved. The present experiments do not distinguish the relative contributions of membrane-associated DDR1 and extracellularly released DDR1-ECD to the observed treatment effects. In addition, collagen remodeling was characterized primarily by histological and SHG imaging, without quantitative assessment of fiber organization or tissue mechanics. Finally, although mouse-level transcriptomic analyses identified reproducible neutrophil pathway changes and Ly6G-directed depletion modified treatment response, the functional contribution of individual neutrophil states remains undefined. Further studies incorporating target-engagement measurements, quantitative matrix characterization, and more selective immune-cell perturbations will help clarify the mechanisms underlying these observations.

## Materials and Methods

### Study design and animal ethics

Animal studies were designed to evaluate the tumor localization, antitumor activity, stromal effects, and immune-associated responses of DDR1 nanobody–Fc proteins in murine breast and pancreatic tumor models. BALB/c nude mice were used for in vivo fluorescence imaging, whereas immunocompetent C57BL/6 mice were used for E0771 mammary tumor and KPC1199 pancreatic tumor studies. Mice were acclimated for 1 week before experimental procedures. Animals were assigned to treatment groups by randomization, and experiments were performed without blinding.

The in vivo imaging study included three mice per group. The discovery-stage E0771 screen included three mice per group, whereas the independent E0771 and subcutaneous KPC1199 candidate-comparison studies included seven mice per group. The orthotopic KPC1199 efficacy study included five mice per group, the single-cell RNA-sequencing study included four mice per group, and the anti-PD-1 and anti-Ly6G combination studies each included six mice per group. Assay-specific sample sizes are indicated in the corresponding figure legends.

All animal procedures were conducted in accordance with institutional guidelines for the care and use of laboratory animals and were approved by the Experimental Animal Ethics Committee of Fudan University School of Pharmacy. Mice were monitored regularly for tumor growth, body condition, and general health, and predefined humane endpoints were applied in accordance with the approved protocol.

### Cell lines and culture conditions

Human MDA-MB-468 and MCF7 breast cancer cells, murine MC38 colon carcinoma cells, murine E0771 mammary carcinoma cells, and murine KPC1199 pancreatic ductal adenocarcinoma cells were kindly provided by the laboratories of Min Huang at the Shanghai Institute of Materia Medica, Chinese Academy of Sciences, and Jun Chen at the School of Pharmacy, Fudan University. KPC1199 cells were originally derived from a murine KPC pancreatic ductal adenocarcinoma model. Cells were maintained in high-glucose Dulbecco’s modified Eagle’s medium (DMEM) supplemented with 10% fetal bovine serum, 100 U/ml penicillin, and 100 μg/ml streptomycin at 37°C in a humidified atmosphere containing 5% CO₂. Cells were routinely passaged during logarithmic growth. For tumor implantation, E0771 and KPC1199 cells were harvested during logarithmic growth, and only cell preparations with viability >98% were used.

### Nanobody discovery and library screening

Alpacas were immunized subcutaneously five times at 7-day intervals with recombinant human and mouse DDR1 extracellular domains (DDR1-ECDs) formulated with adjuvant. Blood was collected 10 days after the final immunization, and serum reactivity against human and mouse DDR1-ECD was evaluated by ELISA. Peripheral-blood mononuclear cells (PBMCs) were isolated, total RNA was extracted and reverse-transcribed, and VHH-encoding sequences were amplified by nested PCR for construction of phage- and yeast-display libraries.

The phage-display library had an estimated capacity of approximately 2.5 × 10⁸ transformants. Four rounds of panning were performed using alternating human and mouse DDR1-ECD, in the sequence human, mouse, human, and mouse, with progressively increased selection stringency. Enrichment during successive rounds of selection was monitored by colony recovery and antigen-binding screening.

A yeast-display VHH library containing approximately 2 × 10⁹ independent clones was subsequently generated. VHH surface display was monitored through V5-tag expression, with approximately 20.1% of the primary library showing detectable V5-positive display. Library enrichment combined magnetic-activated cell sorting (MACS) and fluorescence-activated cell sorting (FACS). Negative magnetic depletion was first used to reduce nonspecific binders, followed by positive enrichment with biotinylated DDR1-ECD. Antigen-binding, V5-positive yeast populations were further enriched by flow sorting. Individual clones were recovered after selection, screened for binding to both human and mouse DDR1-ECD, and sequenced to identify unique cross-reactive VHH candidates.

### Nanobody expression and purification

His-tagged VHHs and VHH-Fc fusion proteins were transiently expressed in 293F cells. Cells were maintained in 293F Hi-Exp medium at 36–38°C in a humidified atmosphere containing 5% CO₂ with orbital shaking at 115–125 rpm using a 50-mm shaking diameter. Transfection was initiated when the cell density reached approximately 3.0 × 10⁶ cells/ml and cell viability was ≥95%. Plasmid DNA and transfection reagent were separately diluted in 293F Hi-Exp medium, each to 5% of the final culture volume, combined, and incubated at room temperature for 10–15 min to allow formation of DNA–transfection reagent complexes. The complexes were then added gradually to the cell cultures with gentle mixing. At 16–22 h after transfection, 293F Hi-Exp feed was added to 5% (v/v), and cultures were returned to the incubator. Glucose concentrations were monitored daily and maintained above 4 g/l during transient expression. Culture supernatants were harvested on day 7 for protein purification.

His-tagged VHHs were purified from clarified culture supernatants by nickel-affinity chromatography using standard immobilized metal-affinity chromatography procedures. VHH-Fc proteins were purified by Protein A affinity chromatography. For Protein A purification, phosphate-buffered saline (PBS) was used as the equilibration and loading buffer, and bound proteins were eluted with 0.1 M glycine-HCl, pH 3.2. Eluted fractions were immediately neutralized with Tris-HCl buffer, pH 8.8, concentrated using 30-kDa molecular-weight-cutoff centrifugal filters, and buffer-exchanged into PBS by repeated concentration and dilution. Purified proteins were assessed by SDS-PAGE before subsequent biochemical and in vivo studies.

### Enzyme-linked immunosorbent assay

Binding of the four prioritized DDR1 nanobody–Fc proteins to recombinant DDR1 extracellular domains was assessed by enzyme-linked immunosorbent assay (ELISA). For cross-species binding analysis, recombinant human DDR1-ECD and mouse DDR1-ECD were immobilized on microplates, and serially diluted DDR1nb-3-Fc, DDR1nb-8-Fc, DDR1nb-13-Fc, or DDR1nb-31-Fc was added to generate concentration–response curves. Bound nanobody–Fc proteins were detected using an HRP-conjugated secondary antibody directed against human IgG, followed by TMB development. Absorbance was measured at 450 nm using a Multiskan FC microplate reader (Thermo Scientific), and EC50 values were estimated by nonlinear regression. Three replicate measurements were performed for each condition.

Preferential recognition of DDR1 over DDR2 was evaluated in a separate ELISA using recombinant human or mouse DDR1-ECD and the corresponding DDR2-ECD proteins. Antigens were diluted in carbonate coating buffer (pH 9.6) to 1 μg/ml and added to MaxiSorp plates at 100 μl per well for overnight coating at 4°C. Plates were washed five times with PBS containing 0.05% Tween 20 (PBST) and blocked with 3% bovine serum albumin in PBS at 37°C for 2 h. DDR1nb-Fc proteins were tested at 10, 2, 0.4, 0.08, and 0.016 μg/ml and incubated at 37°C for 1 h. After five washes with PBST, bound proteins were detected with an HRP-conjugated anti-human IgG antibody. TMB substrate was added at 100 μl per well and incubated for 10 min, after which the reaction was stopped with 50 μl of 2 M HCl. Absorbance was measured at 450 nm. Each condition was measured in triplicate.

### Biolayer interferometry

Binding kinetics of DDR1nb-3-Fc, DDR1nb-8-Fc, DDR1nb-13-Fc, and DDR1nb-31-Fc to recombinant human and mouse DDR1-ECD were measured by biolayer interferometry (BLI). Anti-human IgG Fc capture (AHC) biosensors were loaded with each DDR1nb-Fc protein at 5 μg/ml for 80 s. Measurements were performed in PBS containing 0.02% Tween 20 at 25°C. Recombinant human DDR1-ECD-His and mouse DDR1-ECD-His were used as analytes at 50, 25, 12.5, 6.25, 3.13, and 0 nM. After a 60-s baseline equilibration, association was monitored for 180 s, followed by a 300-s dissociation phase. Sensorgrams were fitted using a Langmuir binding model to estimate the association rate constant (k_on), dissociation rate constant (k_off), and equilibrium dissociation constant (K_D). Because the analyzed ligands were Fc-fused VHH molecules, the resulting kinetic constants are reported as apparent binding parameters for the assay configuration used.

### Cell-associated and tumor-section binding

Cell-associated binding of the four prioritized DDR1nb-Fc proteins was evaluated using murine E0771 cells. To assess the potential influence of cell-detachment procedures on detectable surface DDR1, adherent E0771 cells were collected using trypsin, Accutase, or mechanical scraping. The resulting cell suspensions were incubated with DDR1nb-3-Fc, DDR1nb-8-Fc, DDR1nb-13-Fc, DDR1nb-31-Fc, or human IgG1 Fc control. After washing, cell-associated DDR1nb-Fc was detected using a fluorescently labeled anti-human Fc secondary antibody and analyzed by flow cytometry. Binding was quantified as the percentage of antibody-positive cells among total analyzed cells. Three independent experiments were performed.

Binding to tumor tissue was evaluated using sections of E0771 tumors. Sections were incubated with human IgG1 Fc control or the indicated DDR1nb-Fc proteins, followed by detection with an HRP-conjugated anti-human IgG1 antibody and chromogenic development. Stained sections were imaged to assess tumor-tissue reactivity of the candidate proteins.

### CCK-8 cell-viability assay

MDA-MB-468, MCF7, and MC38 cells were seeded into 96-well plates at 3,000 cells per well and allowed to adhere overnight. The following day, cells were treated with DDR1nb-3-Fc, DDR1nb-8-Fc, DDR1nb-13-Fc, or DDR1nb-31-Fc at final concentrations of 0, 0.1, 1, 10, or 100 μg/ml. After 48 h of treatment, Cell Counting Kit-8 (CCK-8) reagent was added at one-tenth of the culture volume, and cells were incubated for an additional 2 h at 37°C. Absorbance at 450 nm was measured using a Multiskan FC microplate reader (Thermo Scientific).

### Cy7 labeling and in vivo fluorescence imaging

DDR1nb-Fc proteins and human IgG1 Fc control were fluorescently labeled using sulfo-Cy7 NHS ester. Sulfo-Cy7 NHS ester was dissolved in DMSO, and 0.1 M NaHCO₃ was added to the protein solution at 25% of the protein-solution volume. The dye was then added at a 10-fold molar excess relative to protein, and the reaction mixture was incubated with gentle rotation for 1 h at room temperature. Unreacted dye was removed by dialysis. Protein and dye concentrations were determined after labeling, with Cy7 absorbance measured at 740 nm to estimate the dye-to-protein labeling ratio.

For in vivo imaging, E0771 cells (5 × 10⁶ cells in 100 μl PBS) were implanted subcutaneously into the right dorsal flank of BALB/c nude mice. When tumors reached approximately 200 mm³, mice received Cy7-labeled human IgG1 Fc control or the indicated DDR1nb-Fc protein by intraperitoneal injection. Whole-body fluorescence images were acquired at 12, 24, 36, 48, and 72 h after administration using an IVIS Spectrum imaging system (PerkinElmer); n = 3 mice per group. Tumor-associated fluorescence was quantified as the fraction of whole-body fluorescence localized to the tumor region.

### Subcutaneous E0771 and KPC1199 tumor models

Five-week-old C57BL/6 mice were acclimated for 1 week before tumor implantation. Female mice were used for the E0771 mammary tumor model, whereas male mice were used for the KPC1199 pancreatic ductal adenocarcinoma model. E0771 or KPC1199 cells in logarithmic growth were harvested, resuspended in PBS at 5 × 10⁷ cells/ml, and only preparations with cell viability >98% were used for implantation. Each mouse received 5 × 10⁶ tumor cells in 100 μl PBS by subcutaneous injection into the right dorsal flank. Tumor growth was monitored longitudinally by measuring the longest diameter (L) and the perpendicular shorter diameter (W), and tumor volume was estimated as V=0.5×L×W^2^.

For the definitive candidate-comparison studies, treatment was initiated when tumor volume reached at least 80 mm³. Mice received human IgG1 Fc control, DDR1nb-3-Fc, DDR1nb-8-Fc, DDR1nb-13-Fc, or DDR1nb-31-Fc at 10 mg/kg by intraperitoneal injection twice weekly. Seven mice were included per group. Body weight and tumor growth were monitored throughout treatment. E0771 tumors were collected on day 21 after tumor implantation, whereas subcutaneous KPC1199 tumors were collected on day 26. Excised tumors were weighed and processed for histological, collagen, and immune analyses.

For the discovery-stage E0771 screen, ten DDR1nb-Fc candidates were evaluated in the same syngeneic tumor model at 10 mg/kg by intraperitoneal injection twice weekly, with three mice per group. Tumor growth and endpoint tumor weight were used for in vivo candidate prioritization.

### Orthotopic pancreatic tumor model

Five-week-old male C57BL/6 mice were acclimated for 1 week before tumor implantation. KPC1199 cells in logarithmic growth were harvested and resuspended at 2 × 10⁷ cells/ml in a 1:1 mixture of PBS and Matrigel. At approximately 6 weeks of age, mice underwent surgical implantation of KPC1199 cells into the pancreatic tail. The pancreas was exposed through an approximately 1-cm left flank incision, and 50 μl of the cell suspension, corresponding to 1 × 10⁶ KPC1199 cells, was injected into the pancreatic tail.

Seven days after tumor implantation, mice were treated with human IgG1 Fc control or DDR1nb-3-Fc at 10 mg/kg by intraperitoneal injection twice weekly. Five mice were included per group in the orthotopic efficacy study. At the experimental endpoint, pancreatic tumors were excised and weighed, and tumor tissues and major organs were collected for histological and other downstream analyses.

### PD-1 and anti-Ly6G combination experiments

For the E0771 combination study, female C57BL/6 mice were implanted subcutaneously with 5 × 10⁶ E0771 cells in 100 μl PBS as described above. When tumors reached at least 80 mm³, mice were randomly assigned to receive human IgG1 Fc control, anti-PD-1 antibody, DDR1nb-3-Fc, or DDR1nb-3-Fc plus anti-PD-1 antibody. DDR1nb-3-Fc and anti-mouse PD-1 antibody (clone RMP1-14; Bio X Cell) were each administered at 10 mg/kg by intraperitoneal injection twice weekly. Six mice were included per group. Tumor growth was monitored longitudinally, and tumor volume was estimated as V=0.5×L×W^2^. Tumors were collected and weighed on day 19 after tumor implantation.

In an independent subcutaneous KPC1199 experiment, male C57BL/6 mice were assigned to receive human IgG1 Fc control, anti-Ly6G antibody, DDR1nb-3-Fc, or DDR1nb-3-Fc plus anti-Ly6G antibody. Anti-Ly6G antibody (clone 1A8; TargetMol) was administered intraperitoneally at 12.5 μg per mouse in 100 μl PBS once daily, beginning on day 4 after tumor inoculation. DDR1nb-3-Fc was administered intraperitoneally at 10 mg/kg twice weekly. Six mice were included per group. Tumor growth was monitored through day 32, when tumors were excised and weighed. Peripheral blood and major organs were collected at the same endpoint for hematologic and organ-weight analyses.

### Histology and Masson’s trichrome staining

Tumor tissues and major organs collected at the indicated experimental endpoints were fixed in 4% paraformaldehyde, dehydrated through graded ethanol, and embedded in paraffin. Paraffin blocks were sectioned at approximately 4–5 μm. For hematoxylin and eosin (H&E) staining, sections were deparaffinized, rehydrated, stained using standard H&E procedures, dehydrated, cleared, and mounted for histopathological evaluation.

For Masson’s trichrome staining, paraffin sections were deparaffinized in xylene and rehydrated through graded ethanol. Sections were stained using a Masson’s trichrome staining procedure to visualize collagen-containing stromal regions and tissue morphology, followed by dehydration and mounting. Whole-slide images were acquired using an Olympus VS200 slide-scanning system. Masson-stained sections were evaluated for the distribution and organization of collagen-rich regions within tumor tissues.

### Second-harmonic generation imaging

Freshly collected tumor tissues were embedded in optimal cutting temperature (OCT) compound and stored at −80°C until sectioning. Before cryosectioning, OCT-embedded tissues were equilibrated at −20°C for at least 2 h and sectioned at 20 μm using a cryostat. Sections were allowed to equilibrate at 37°C for 30 min and counterstained with 5 μM DCS1. After mounting with Fluoromount-G antifade medium, sections were imaged using a Leica TCS SP8 multiphoton microscope equipped with a 20× HC PL Apo objective (NA 0.7).

Collagen-associated second-harmonic generation (SHG) signals were excited at 840 nm and collected using a 420 ± 5 nm detection window. Imaging focused on tumor-border regions to visualize the organization and continuity of collagen-rich structures at the tumor margin. SHG images were used to assess treatment-associated changes in collagen architecture rather than absolute collagen abundance.

### Immunohistochemistry and dual chromogenic staining

Tumor tissues were fixed in 4% paraformaldehyde overnight at 4°C, dehydrated through graded ethanol, embedded in paraffin, and sectioned at 4–5 μm. Sections were mounted on pretreated glass slides and dried at 60°C for 1 h. After deparaffinization and antigen retrieval, sections were washed three times with PBS for 5 min each and incubated with primary antibodies against CD45, CD8, NK1.1, or Ki67 overnight at 4°C. After three additional PBS washes, appropriate secondary antibodies were applied for 1 h at room temperature. Immunoreactivity was visualized by chromogenic development, including DAB where applicable, and sections were subsequently mounted for imaging.

CD45, CD8, and NK1.1 staining in E0771 tumors and Ki67 staining in subcutaneous KPC1199 tumors were quantified using H-scores. For dual chromogenic analysis, E0771 tumor sections were stained for Ly6G and CD62L, with Ly6G-positive staining visualized in brown and CD62L-positive staining in pink. Dual-stained sections were used to assess the spatial distribution of Ly6G- and CD62L-associated signals within the tumor tissue, including inflammatory and necrotic regions.

### Flow-cytometric immune profiling

Tumors and spleens collected at the indicated experimental endpoints were processed into single-cell suspensions for flow-cytometric analysis. For lymphoid immune profiling, cells were stained with fluorophore-conjugated antibodies against CD45, CD3, CD4, CD8, and NK1.1, as appropriate for each experiment. For analysis of the Ly6G-positive myeloid compartment, cells were stained with antibodies against CD45, CD11b, and Ly6G. Samples were analyzed using a CytoFLEX flow cytometer (Beckman Coulter).

Intratumoral CD45-positive, CD3-positive, CD4-positive, CD8-positive, and NK1.1-positive populations were quantified as percentages of total analyzed tumor cells. In the E0771 model, CD45-positive CD11b-positive Ly6G-positive cells were quantified both as a percentage of total analyzed tumor cells and as a percentage of the parental CD45-positive population. Corresponding Ly6G-positive populations in spleens were quantified relative to total analyzed splenic cells and to the parental CD45-positive compartment, as indicated. Flow-cytometry gates were defined consistently across treatment groups within each experiment.

### Serum enzymes and inflammatory-marker assays

Blood was collected from tumor-bearing mice at the indicated experimental endpoints, and serum was isolated for biochemical and inflammatory-marker analyses. In the E0771 study, serum aspartate aminotransferase (AST) and alanine aminotransferase (ALT) concentrations were measured as short-term indicators of hepatocellular injury. Serum concentrations of IL-6, IFN-γ, TNF-α, and IL-1β were also determined at the study endpoint. In parallel, tumor-associated TNF-α and IL-1β levels were measured separately from the corresponding E0771 tumors. In the subcutaneous KPC1199 candidate-comparison study, endpoint serum TNF-α and IL-1β concentrations were measured to assess treatment-associated changes in circulating inflammatory mediators. Assay-specific sample sizes are indicated in the corresponding figure legends.

### Single-cell sample preparation, library construction, and upstream processing

Orthotopic KPC1199 tumors were collected from four human IgG1 Fc control mice and four DDR1nb-3-Fc-treated mice. Adipose, fibrous, and visibly necrotic tissues were removed, and the remaining tumor tissue was cut into approximately 2–4-mm fragments. Tumors were dissociated using a Tumor Dissociation Kit in gentle MACS C Tubes, followed by incubation at 37°C for 40 min and mechanical dissociation using a gentle MACS Dissociator. The resulting cell suspensions were passed through 70-μm MACS Smart Strainers to remove residual tissue debris. CD45-positive tumor-associated immune cells were subsequently enriched by positive magnetic selection using the EasySep Mouse TIL (CD45) Positive Selection Kit, washed, and resuspended for single-cell library preparation.

Single-cell libraries were generated using the DNBelab C4 platform and sequenced in paired-end mode. The sequencing-provider report described quality-control criteria of 200–10,000 detected genes per cell and a mitochondrial transcript fraction of ≤20%, together with DoubletFinder-based doublet prediction. The downstream analyses in this study were performed using the recovered annotated Seurat objects. These objects retained a subset of cells labeled as predicted doublets in the provider’s pre-filtering metadata, and the historical decisions underlying doublet exclusion could not be fully reconstructed.

The recovered all-cell Seurat object contained 57,907 cells. Its saved processing records indicated LogNormalize normalization with a scaling factor of 10,000, principal-component analysis using 20 components, and clustering at a resolution of 0.6. The recovered neutrophil subset contained 22,923 cells, with subsequent Harmony-based analyses using dimensions 1–20 and a clustering resolution of 0.4. Cell-type and neutrophil-state annotations were evaluated using marker-expression profiles. The original filtering decisions underlying the construction of the neutrophil subset could not be independently recovered.

### Single-cell visualization and mouse-level compositional analysis

Downstream analyses for the present study were performed using the recovered processed single-cell objects, including the existing cell annotations and UMAP coordinates. The final CD45-enriched dataset contained 57,907 cells assigned to nine major immune-cell classes. The neutrophil compartment contained 22,923 cells and was resolved into seven transcriptional states: Neu-Gpr84, Neu-Prss2, Neu-Ero1a, Neu-Chil3, Neu-Ifit1, Neu-Lrg1, and Neu-Ext1. Cell-class and neutrophil-state annotations were evaluated using their corresponding marker-expression profiles.

For marker-expression dot plots, dot size represents the percentage of cells with detected expression of the indicated gene, whereas dot color represents the gene-wise z-scored mean log-normalized expression across the displayed cell classes or neutrophil states. Group-split UMAP visualizations used equal-sized cell samples for display only; all retained cells from all eight mice were used for quantitative abundance analyses.

Cell-class frequencies were calculated separately for each mouse as the proportion of recovered cells assigned to each major immune-cell class. Neutrophil-state frequencies were calculated separately for each mouse within that mouse’s neutrophil compartment. Treatment effects were expressed as DDR1nb-3-Fc-minus-control differences in percentage points. Mice, rather than individual cells, were treated as the independent experimental units. Group differences were evaluated using two-sided Welch’s t-tests, and P values were adjusted using the Benjamini–Hochberg procedure separately across the nine major cell-class comparisons and the seven neutrophil-state comparisons. Adjusted P values are reported as q values, with q < 0.05 considered statistically supported after multiple-testing correction. Error bars for treatment-minus-control differences represent nominal 95% Welch confidence intervals.

### Mouse-level neutrophil pseudobulk and pathway analyses

Raw RNA counts from the 22,923 annotated neutrophils were aggregated within each of the eight mice to generate mouse-level pseudobulk libraries, with mice treated as the independent biological replicates. Differential expression was analyzed using edgeR v4.4.0 after low-expression filtering with filterByExpr, TMM normalization, and quasi-likelihood negative-binomial modeling with treatment as the design factor; P values were adjusted using the Benjamini–Hochberg procedure. Genes were ranked by sign (log2 fold change) × sqrt (quasi-likelihood F statistic) for pre-ranked GSEA using GSEApy v1.3.1 against the MSigDB Hallmark, Reactome, Gene Ontology Biological Process, and KEGG Mouse collections, with gene-set P values adjusted separately within each collection. Neutrophil-state composition was additionally evaluated at the mouse level using centered log-ratio transformation and Aitchison-distance-based exact PERMANOVA across all 70 possible 4-versus-4 treatment-label assignments. State-resolved pseudobulk/GSEA and leave-one-mouse-out analyses were performed to assess whether pathway-level effects were confined to individual neutrophil states or driven by a single mouse.

### Statistical analysis

Data are presented as mean ± SD unless otherwise indicated. Statistical analyses of conventional experimental data were performed using GraphPad Prism 8 (GraphPad Software). Comparisons between two independent groups were performed using an unpaired two-tailed t test. Comparisons among multiple treatment groups at a single endpoint were generally performed using one-way analysis of variance (ANOVA), whereas two-way ANOVA was used for experiments involving two factors or longitudinal measurements, as specified in the corresponding figure legends. For the anti-Ly6G co-treatment experiment, endpoint tumor weights were analyzed using two-way factorial ANOVA with DDR1nb-3-Fc treatment and anti-Ly6G treatment as the two factors. The DDR1nb-3-Fc × anti-Ly6G interaction term was used to test whether Ly6G-positive cell depletion modified the effect of DDR1nb-3-Fc treatment. For conventional experimental analyses, p < 0.05 was considered statistically significant. Exact sample sizes and the statistical tests applied to individual experiments are provided in the corresponding figure legends.

## Resource availability

### Lead contact

Further information and requests for resources and reagents should be directed to and will be fulfilled by the lead contact, Li Ye.

### Materials availability

DDR1 nanobody–Fc proteins and their corresponding expression constructs were generated in this study. Requests for these materials should be directed to the lead contact. Material availability is subject to any applicable intellectual property or material transfer restrictions, as specified by the relevant rights holders.

### Data and code availability

- Data: The processed single-cell RNA-sequencing data generated in this study have been submitted to the Open Archive for Miscellaneous Data (OMIX) of the National Genomics Data Center (https://ngdc.cncb.ac.cn/omix) under accession number OMIX021906. Pending public release of the repository record, access to the processed datasets may be requested from the lead contact. Raw sequencing reads are being retrieved from the sequencing provider and have not yet been deposited in a public sequencing archive. Additional data supporting the findings of this study are available from the lead contact upon reasonable request.
- Code: This study does not report original code.
- Any additional information required to reanalyze the data reported in this paper is available from the lead contact upon request.

## Acknowledgments

This work was supported by the Macao Science and Technology Development Fund (No. FDCT 0019/2024/RIA1, FDCT 0148/2022/A3 and 0013/2026/RIB1) and the Macao Young Scholars Program (AM2025028).

## Author contributions

Conceptualization: X.H., X.L., L.H., and L.Y. Methodology: X.H., X.L., L.H., G.J., and K.K. Investigation: X.H., L.H., G.J., K.K., M.C., Q.Y., M.Y., and S.Y. Formal analysis: X.H., X.L., L.H., M.C., and Q.Y. Visualization: X.H., X.L., and L.H. Resources: Y.Z.Z. and L.Y. Funding acquisition: X.L., Y.Z.Z., and L.Y. Project administration: X.L., Y.Z.Z., and L.Y. Supervision: X.L., Y.Z.Z., and L.Y. Writing—original draft: X.H., X.L., and L.H. Writing—review and editing: X.H., X.L., L.H., G.J., K.K., M.C., Q.Y., M.Y., S.Y., Y.Z.Z., and L.Y.

