## Supplementary Materials for "Extracellular DDR1 targeting remodels collagen-rich tumor stroma and local immunity"

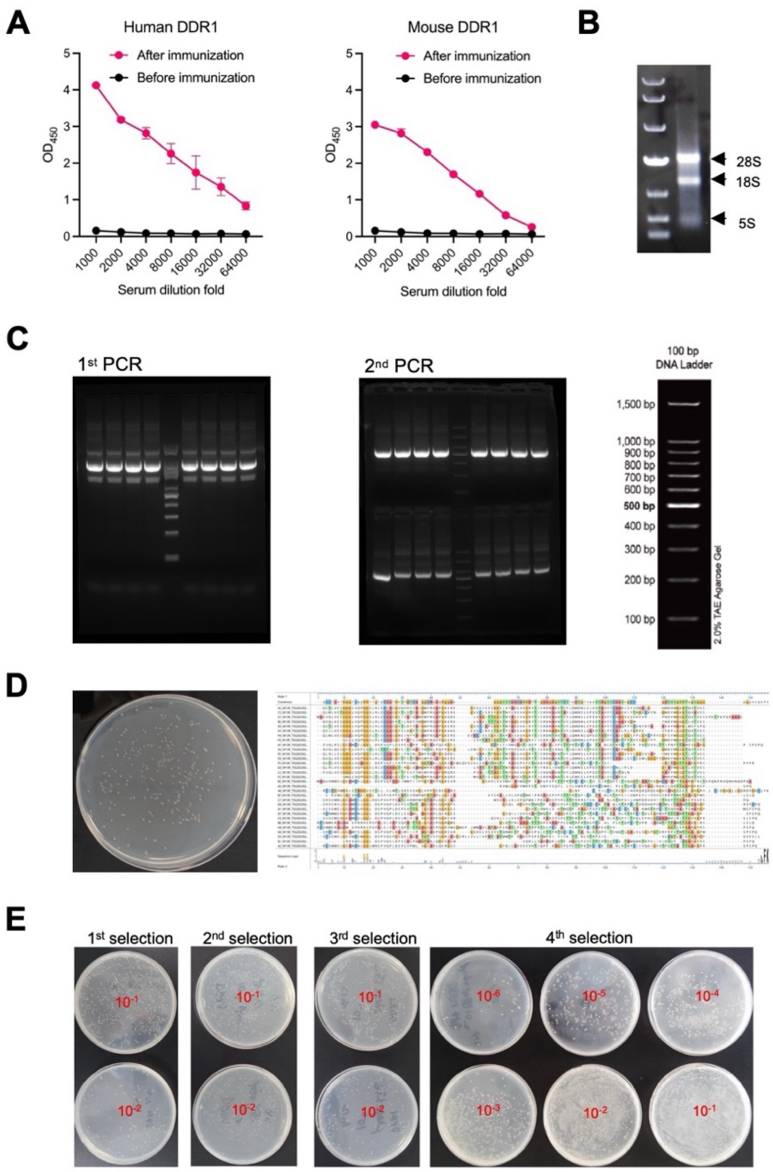


**Supplementary Fig. S1. Construction and enrichment of the phage-display VHH library.**

(A) Serum reactivity against recombinant human and mouse DDR1 extracellular domains (DDR1-ECD) after five rounds of alpaca immunization. (B) Total RNA isolated from alpaca peripheral-blood mononuclear cells (PBMCs). (C) Agarose-gel analysis of PCR-amplified VHH-encoding fragments. (D) Representative transformation plate used to estimate phage-display library size and representative sequence analysis of library clones. The estimated library capacity was $2.5 \times 10^8$ transformants. (E) Representative plates showing enrichment across four rounds of phage-display selection against DDR1-ECD.

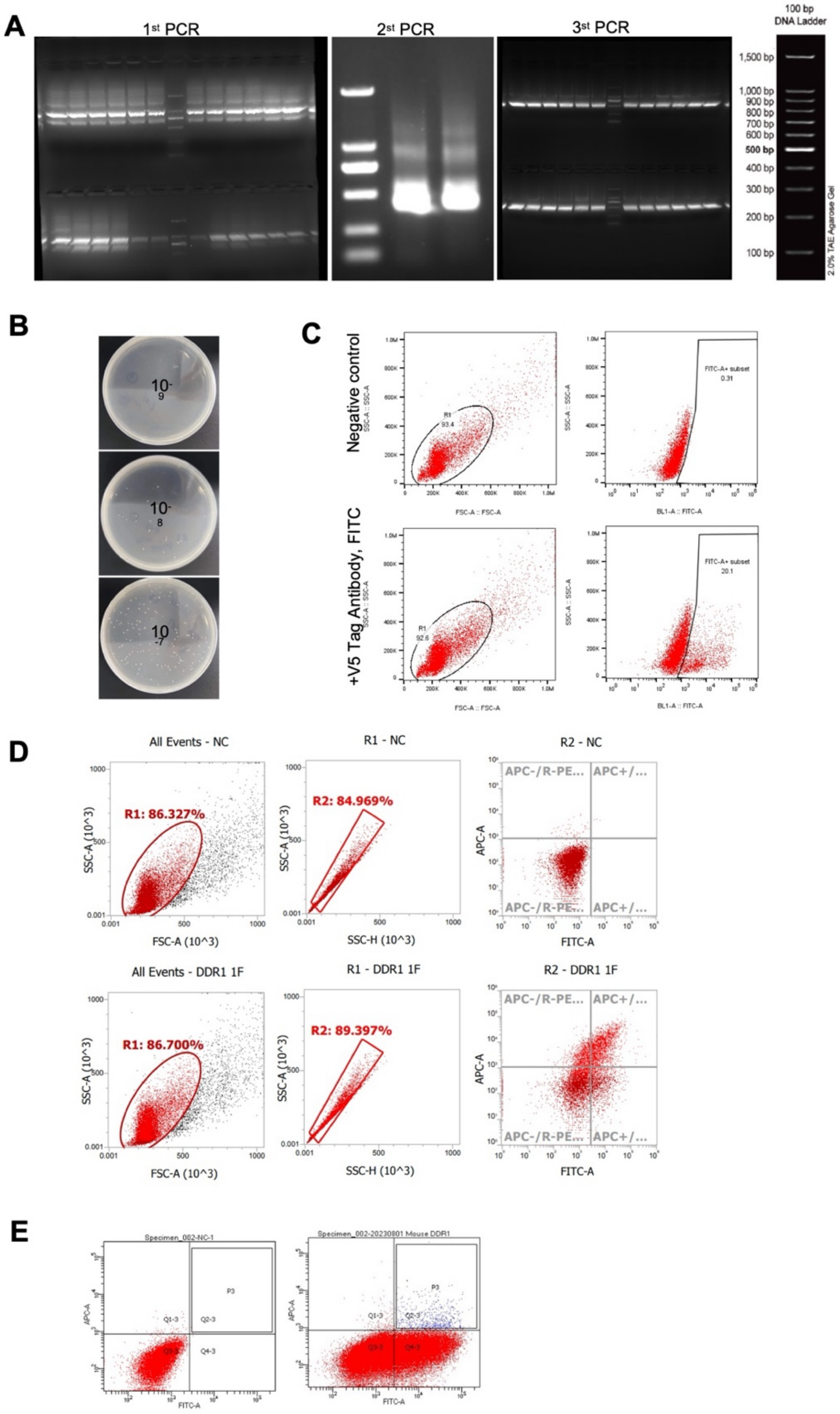


**Supplementary Fig. S2. Construction and enrichment of the yeast-display VHH library.**

(A) Agarose-gel analysis of VHH products generated through three rounds of nested touchdown PCR. The predominant products were approximately 700 base pairs after the first round and approximately 500 base pairs after the second and third rounds. (B) Representative SDCAA plates used to estimate the size of the primary yeast-display library. The estimated library size was approximately 2 × $10^9$ independent clones. (C) Flow-cytometric analysis of V5-tag expression in the primary yeast-display library, showing a display-positive population of 20.1%. (D) Representative flow-cytometry plots of negative-control yeast and yeast cells after the first round of magnetic-activated cell sorting (MACS), showing enrichment of the antigen-binding population. (E) Representative flow-cytometry plots showing gate definition using negative-control yeast and the sorting strategy for the second round of fluorescence-activated cell sorting (FACS).

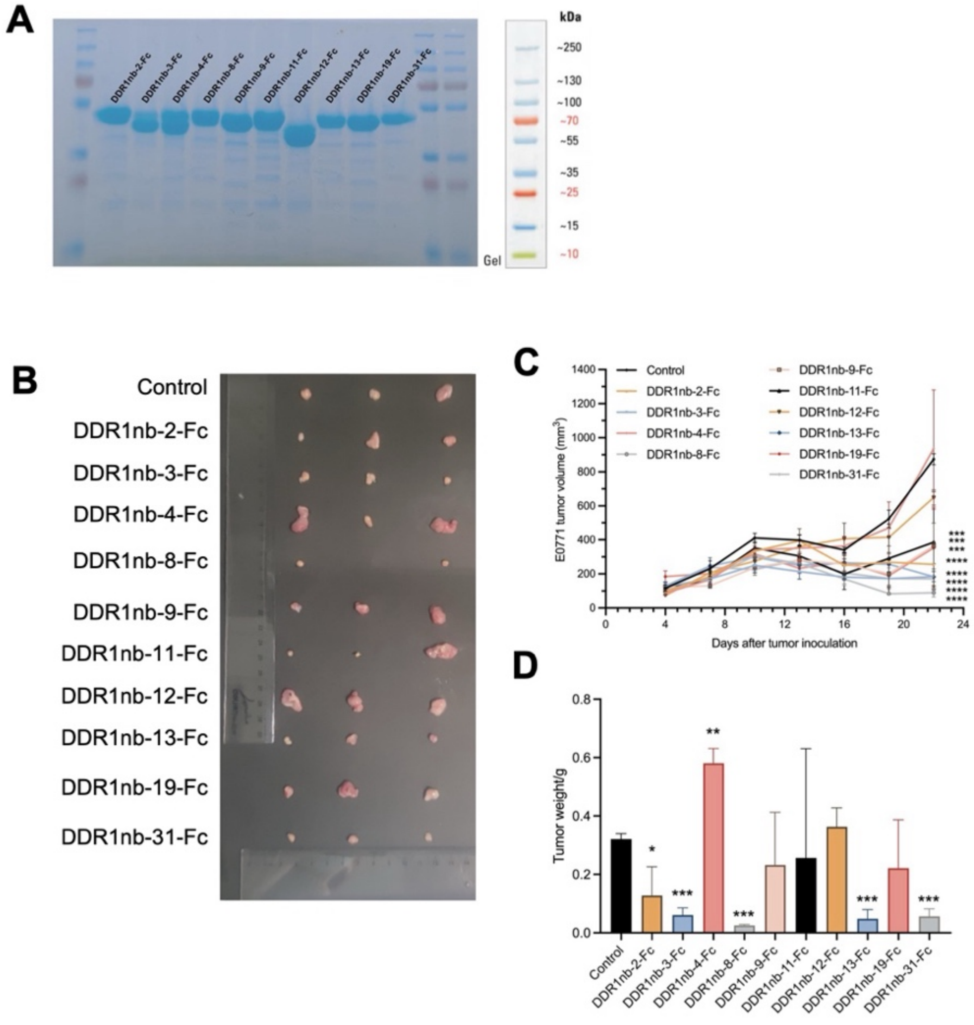


**Supplementary Fig. S3. Discovery-stage in vivo screening of ten DDR1 nanobody–Fc candidates in the E0771 model.**

(A) SDS-PAGE analysis of the ten purified DDR1 nanobody–Fc candidates before in vivo screening. (B) Representative images of excised E0771 tumors collected at the study endpoint from mice receiving human IgG1 Fc control or the indicated DDR1nb-Fc proteins. (C) Longitudinal tumor-volume measurements during the discovery-stage screen. (D) Quantification of endpoint tumor weights. Data are presented as mean ± SD; n = 3 mice per group. Statistical comparisons were performed using one-way ANOVA or two-way ANOVA. Asterisks indicate significance versus the Fc control: *p < 0.05, **p < 0.01, ***p < 0.001, ****p < 0.0001.

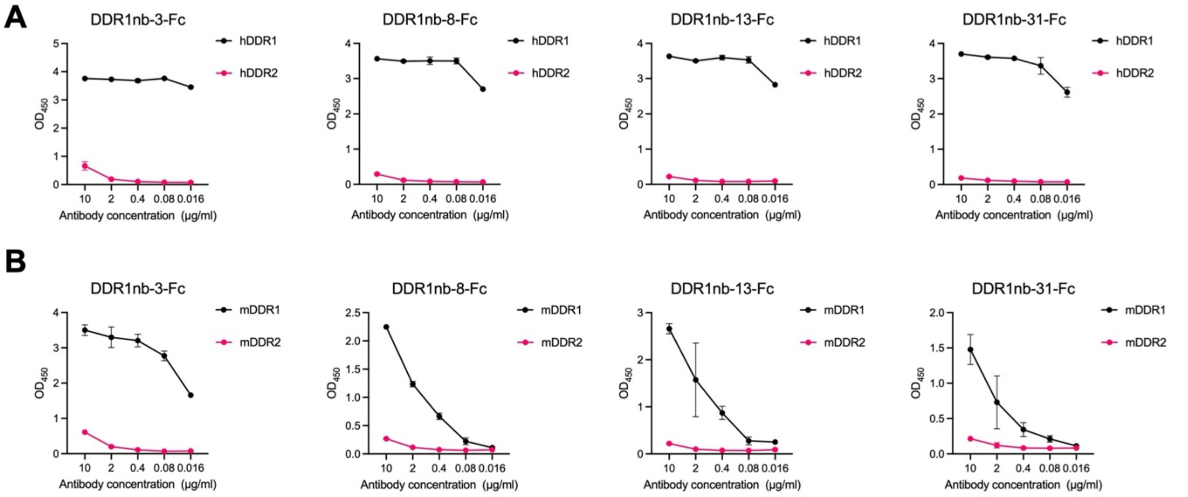


**Supplementary Fig. S4. Preferential binding of DDR1 nanobody–Fc candidates to DDR1 over DDR2 extracellular domains.**

(A) ELISA binding of DDR1nb-3-Fc, DDR1nb-8-Fc, DDR1nb-13-Fc, and DDR1nb-31-Fc to recombinant human DDR1-ECD and DDR2-ECD across the indicated antibody concentrations. (B) ELISA binding of the same DDR1nb-Fc candidates to recombinant mouse DDR1-ECD and DDR2-ECD across the indicated antibody concentrations. Data are presented as mean ± SD from three replicate measurements.

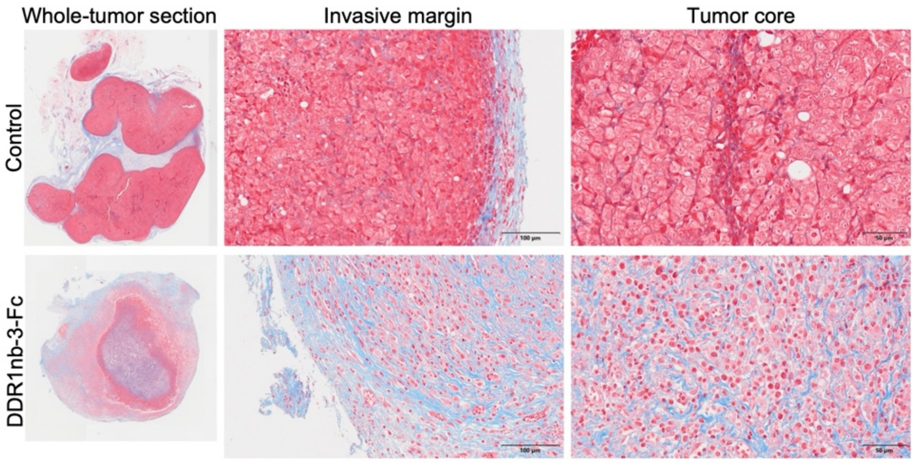


**Supplementary Fig. S5. Regional collagen architecture in E0771 tumors following DDR1nb-3-Fc treatment.**

Representative Masson’s trichrome-stained sections of E0771 tumors from control and DDR1nb-3-Fc-treated mice, shown at the whole-tumor level and at higher magnification in the tumor margin and tumor interior. The images illustrate regional differences in collagen distribution and organization associated with DDR1nb-3-Fc treatment.

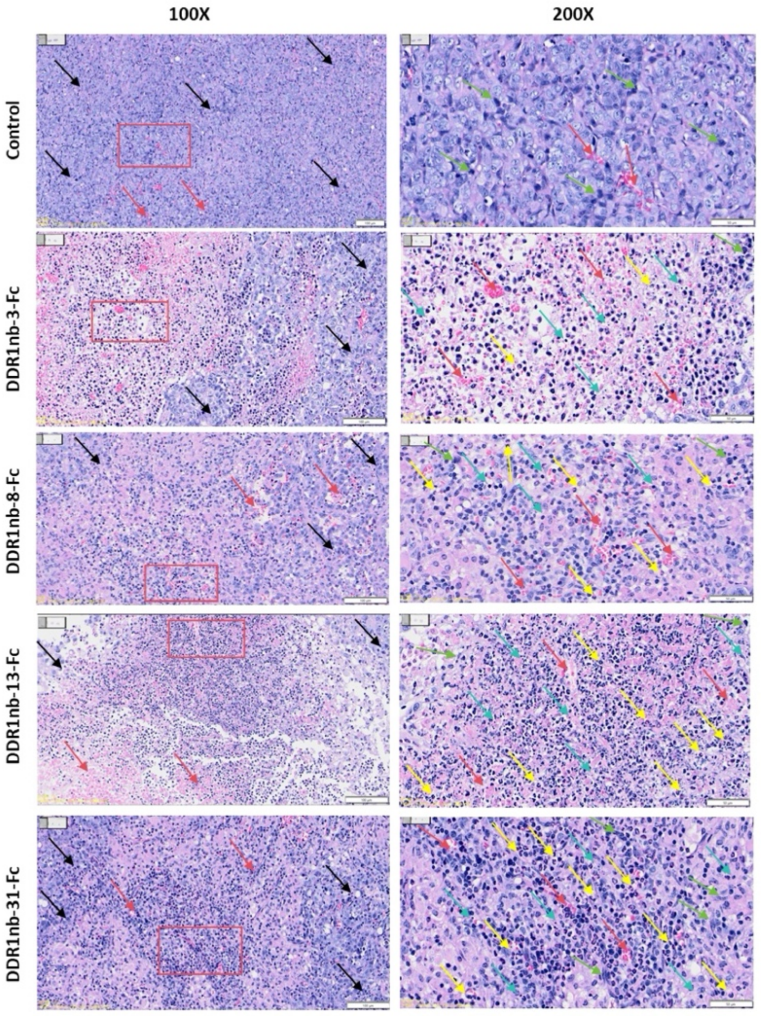


**Supplementary Fig. S6. Histopathological features of E0771 tumors following DDR1 nanobody–Fc treatment.**

Representative hematoxylin-and-eosin (H&E)-stained sections of E0771 tumors from mice receiving human IgG1 Fc control or the indicated DDR1nb-Fc proteins. Images are shown at two magnifications to illustrate tumor morphology and selected histopathological features. Arrows indicate tubular structures (black), pathological mitotic figures (green), hemorrhagic regions (red), lymphocytes (blue), and neutrophils (yellow). Scale bars, 100 μm.

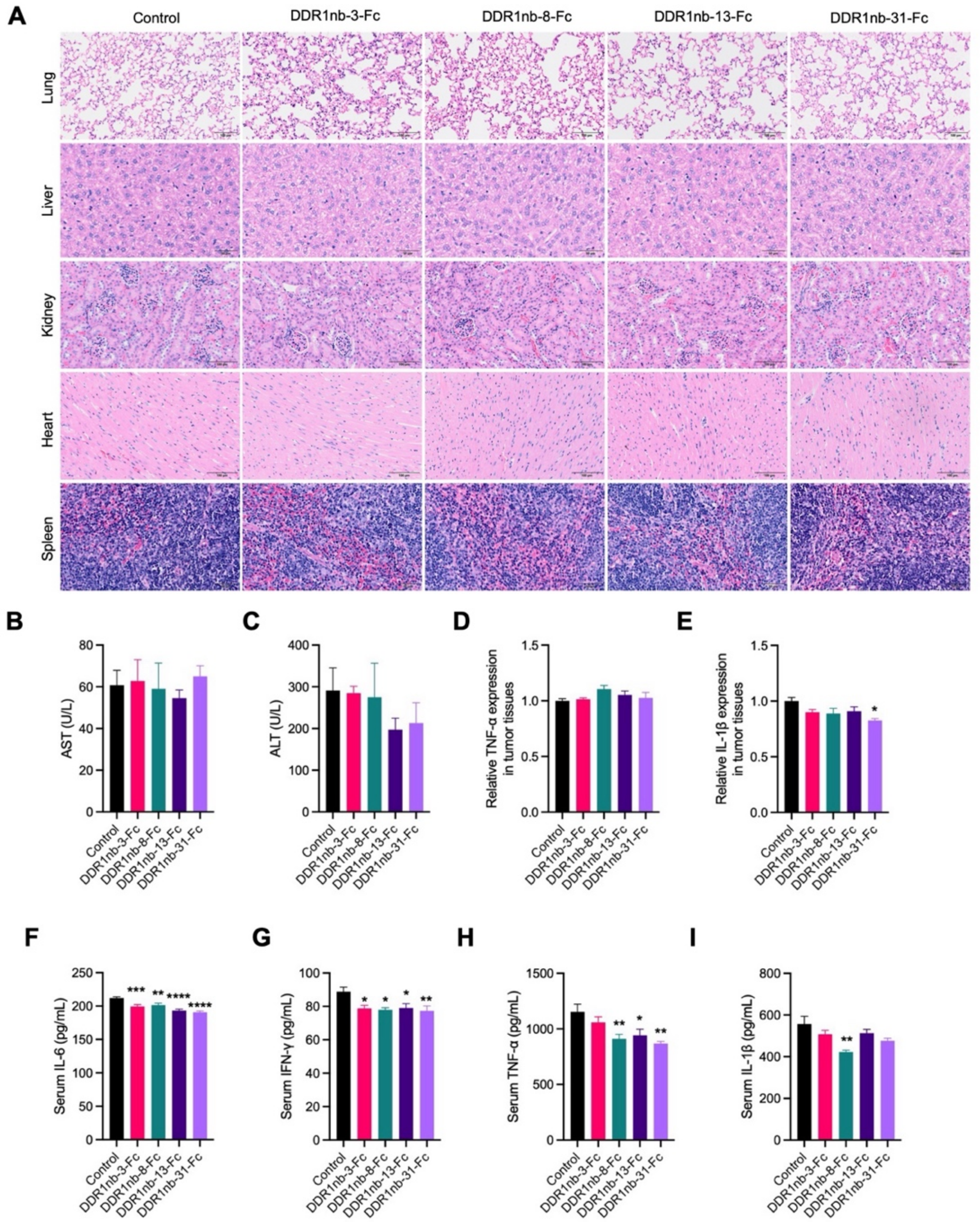


**Supplementary Fig. S7. Endpoint organ histology, serum enzymes, and inflammatory markers in E0771 tumor-bearing mice.**

(A) Representative hematoxylin-and-eosin (H&E)-stained sections of the heart, liver, spleen, lung, and kidney collected at the study endpoint from mice receiving human IgG1 Fc control or the indicated DDR1nb-Fc proteins. (B, C) Serum aspartate aminotransferase (AST) (B) and alanine aminotransferase (ALT) (C) concentrations at the study endpoint. (D, E) Tumor-associated TNF-α (D) and IL-1β (E) concentrations. (F–I) Serum concentrations of IL-6 (F), IFN-γ (G), TNF-α (H), and IL-1β (I). Data are presented as mean ± SD; n = 4–7 mice per group, depending on the assay. Statistical comparisons were performed using one-way ANOVA. Asterisks indicate significance versus the Fc control: *$p < 0.05$, **$p < 0.01$, ***$p < 0.001$, ****$p < 0.0001$.

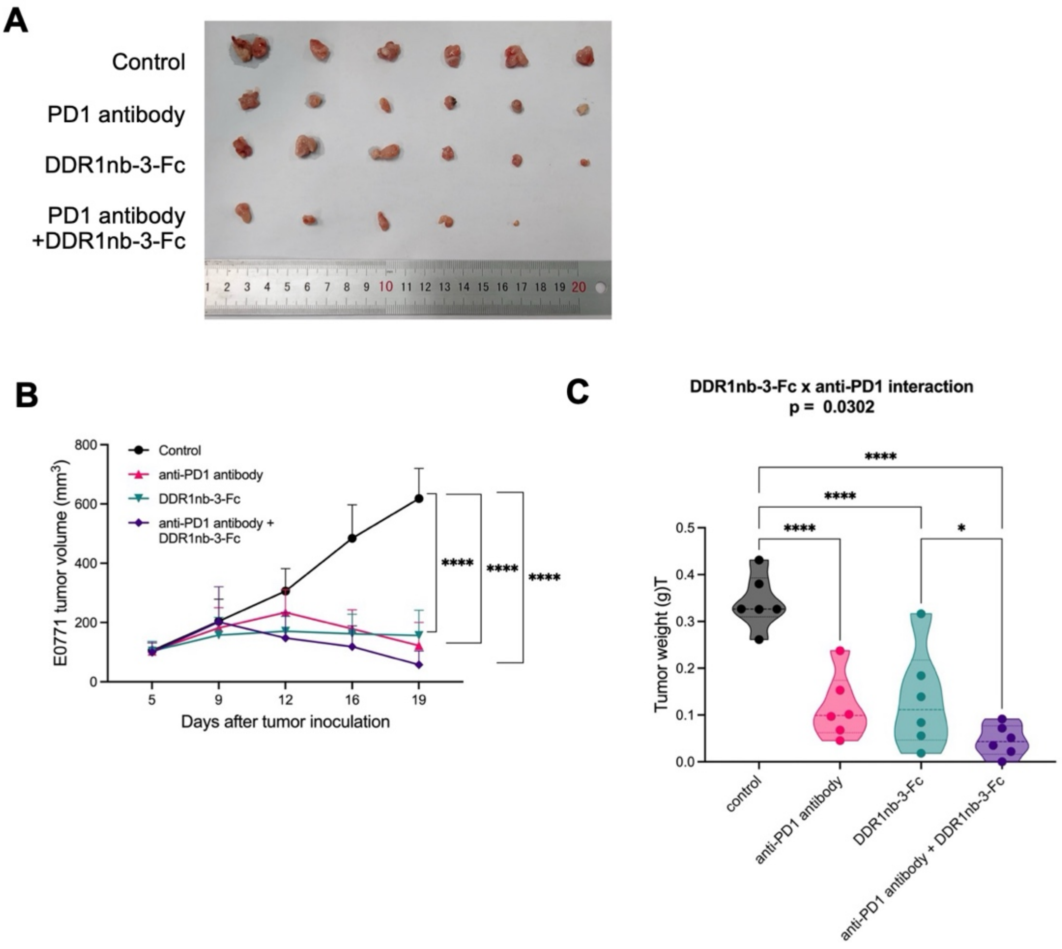


**Supplementary Fig. S8. DDR1nb-3-Fc in combination with PD-1 blockade in E0771 tumors.**

(A) Representative images of excised E0771 tumors collected at the study endpoint from mice receiving control, anti-PD-1 antibody, DDR1nb-3-Fc, or the combination of DDR1nb-3-Fc and anti-PD-1. (B) Longitudinal tumor-volume measurements during treatment. (C) Endpoint tumor weights. Points represent individual mice; n = 6 mice per group. Data are presented as mean ± SD. Longitudinal tumor-volume measurements were analyzed using two-way ANOVA as indicated. Endpoint tumor weights were analyzed using two-way factorial ANOVA with DDR1nb-3-Fc treatment and anti-PD-1 treatment as the two factors. Significant main effects were detected for DDR1nb-3-Fc ($p = 0.0001$) and anti-PD-1 treatment ($p < 0.0001$), together with a significant DDR1nb-3-Fc × anti-PD-1 interaction ($p = 0.0302$). The indicated pairwise comparisons in (C) were evaluated using multiple-comparisons test. Asterisks indicate adjusted P values for the indicated pairwise comparisons: *$p < 0.05$, **$p < 0.01$, ***$p < 0.001$, ****$p < 0.0001$.

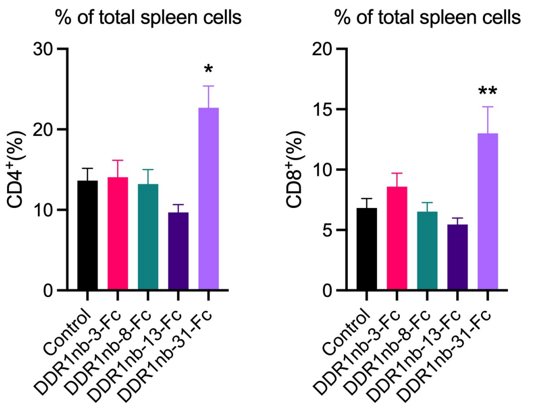


**Supplementary Fig. S9. Splenic CD4-positive and CD8-positive cell frequencies in E0771 tumor-bearing mice following DDR1 nanobody–Fc treatment.**

Flow-cytometric quantification of CD4-positive and CD8-positive cells in spleens collected from E0771 tumor-bearing mice receiving human IgG1 Fc control or the indicated DDR1nb-Fc proteins. Frequencies are expressed as percentages of total analyzed spleen cells. Data are presented as mean ± SD; n = 5 mice per group. Statistical comparisons were performed using one-way ANOVA. Asterisks indicate significance versus the Fc control: *$p < 0.05$, **$p < 0.01$, ***$p < 0.001$, ****$p < 0.0001$.

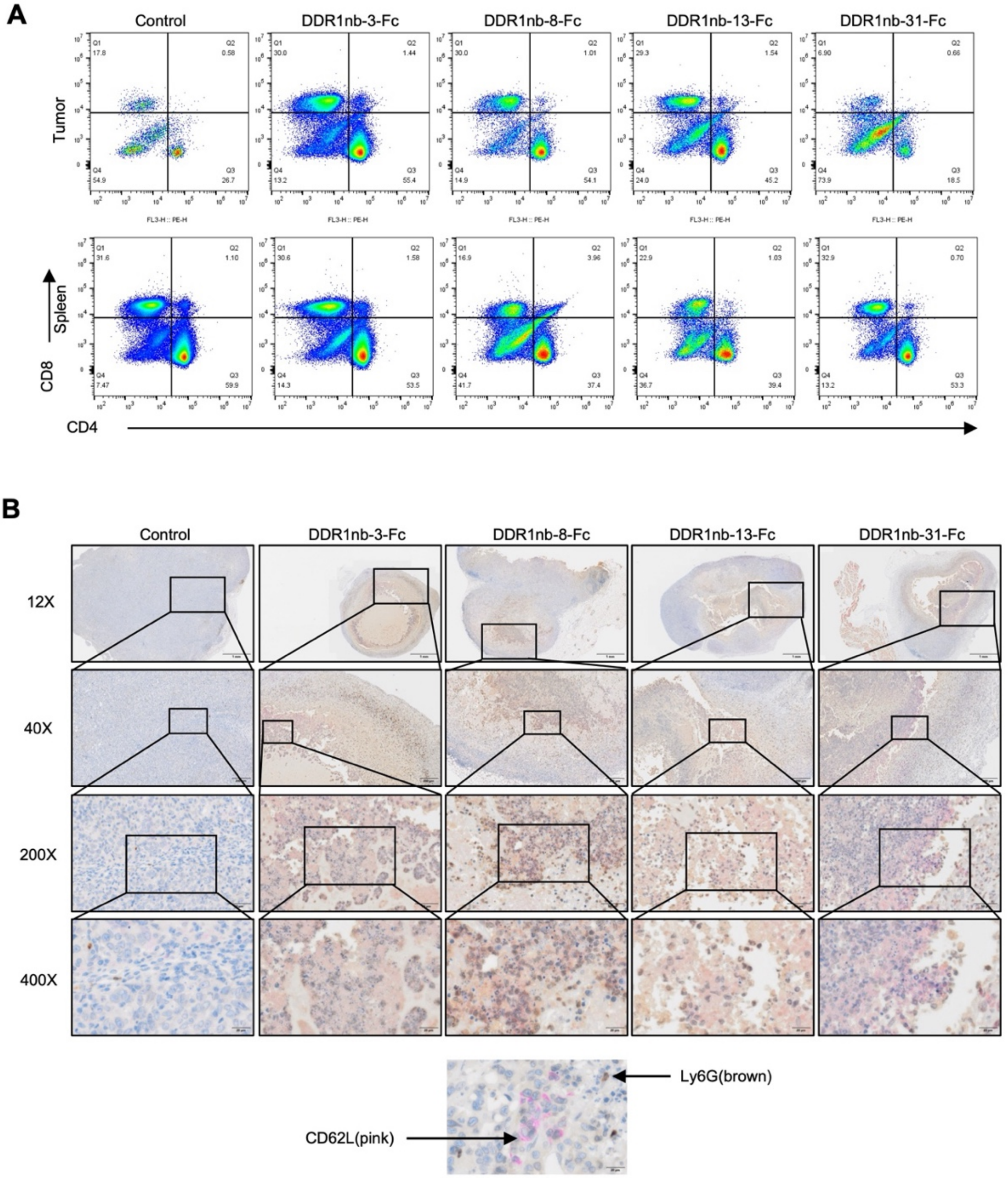


**Supplementary Fig. S10. Representative lymphoid-cell flow-cytometry plots and Ly6G/CD62L staining in the E0771 model.**

(A) Representative CD4-versus-CD8 flow-cytometry plots from E0771 tumors (upper row) and spleens (lower row) collected from mice receiving human IgG1 Fc control or the indicated DDR1nb-Fc proteins. Corresponding quantitative analyses are shown in Fig. 4A and Fig. S9. (B) Representative dual chromogenic staining for Ly6G (brown) and CD62L (pink) in E0771 tumor sections from the indicated treatment groups. Boxed regions indicate areas shown at higher magnification. The bottom inset illustrates the individual chromogenic signals used to identify Ly6G- and CD62L-positive staining. Scale bars, 20 μm.

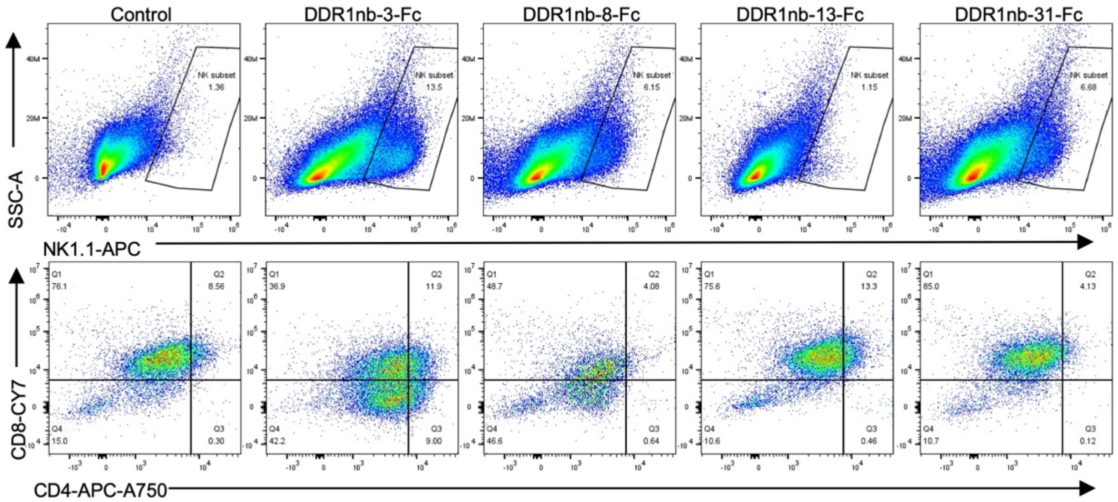


**Supplementary Fig. S11. Representative immune-cell flow-cytometry plots from subcutaneous KPC1199 tumors.**

Representative flow-cytometry plots from subcutaneous KPC1199 tumors collected from mice receiving human IgG1 Fc control or the indicated DDR1nb-Fc proteins. The upper row shows NK1.1-versus-side-scatter profiles, and the lower row shows CD4-versus-CD8 profiles. Corresponding quantitative analyses of intratumoral immune-cell populations are shown in Fig. 5D.

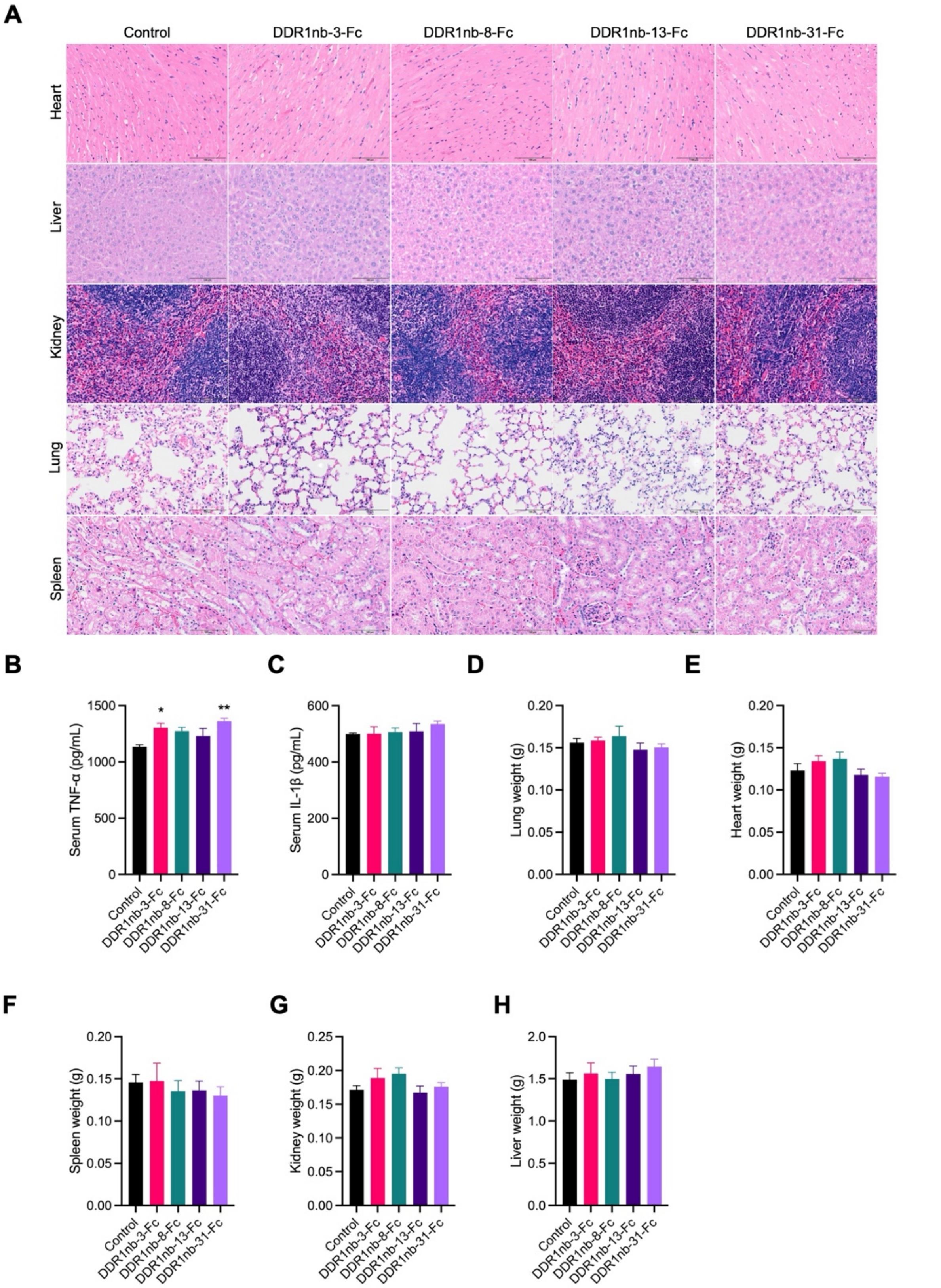


**Supplementary Fig. S12. Endpoint organ histology, inflammatory markers, and organ weights in the subcutaneous KPC1199 model.**

(A) Representative hematoxylin-and-eosin (H&E)-stained sections of the heart, liver, spleen, lung, and kidney collected at the study endpoint from mice receiving human IgG1 Fc control or the indicated DDR1nb-Fc proteins. (B, C) Serum concentrations of TNF-α (B) and IL-1β (C) at the study endpoint. (D–H) Endpoint wet weights of the heart (D), liver (E), spleen (F), lung (G), and kidney (H). Data are presented as mean ± SD; n = 7 mice per group. Statistical comparisons were performed using one-way ANOVA. *$p < 0.05$, **$p < 0.01$, ***$p < 0.001$, ****$p < 0.0001$.

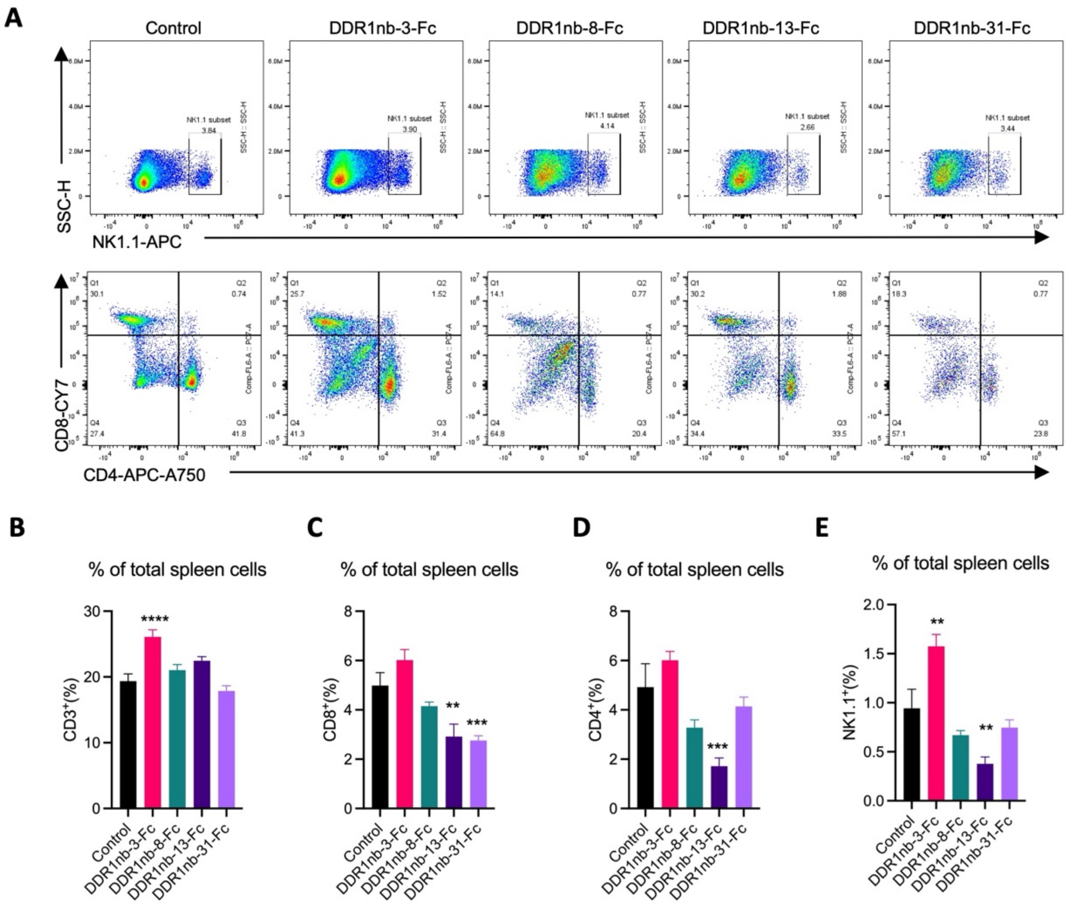


**Supplementary Fig. S13. Splenic lymphoid-cell populations in mice bearing subcutaneous KPC1199 tumors.**

(A) Representative flow-cytometry plots from spleens collected from mice receiving human IgG1 Fc control or the indicated DDR1nb-Fc proteins. The upper row shows NK1.1-versus-side-scatter profiles, and the lower row shows CD4-versus-CD8 profiles. (B–E) Flow-cytometric quantification of CD3-positive (B), CD8-positive (C), CD4-positive (D), and NK1.1-positive (E) cells. Frequencies are expressed as percentages of total analyzed spleen cells. Data are presented as mean ± SD; n = 7 mice per group. Statistical comparisons were performed using one-way ANOVA. Asterisks indicate significance versus the Fc control: *$p < 0.05$, **$p < 0.01$, ***$p < 0.001$, ****$p < 0.0001$.

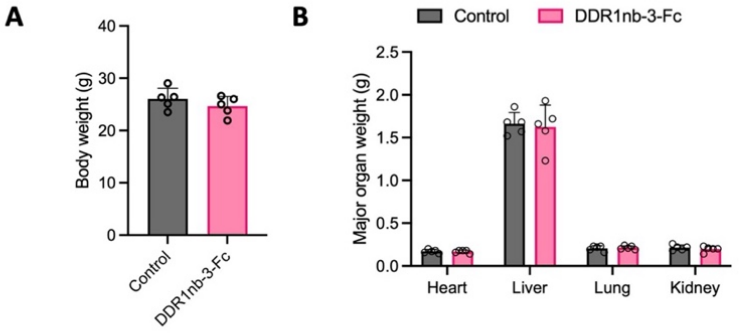


**Supplementary Fig. S14. Endpoint body and organ weights in the orthotopic KPC1199 model.**

(A) Body weights at the study endpoint in mice bearing orthotopic KPC1199 tumors and receiving human IgG1 Fc control or DDR1nb-3-Fc. (B) Endpoint wet weights of the heart, liver, lung, and kidney from the same mice. Points represent individual mice, and bars and error bars indicate mean ± SD; n = 5 mice per group. Statistical comparisons were performed using an unpaired t test for (A) and two-way ANOVA for (B).

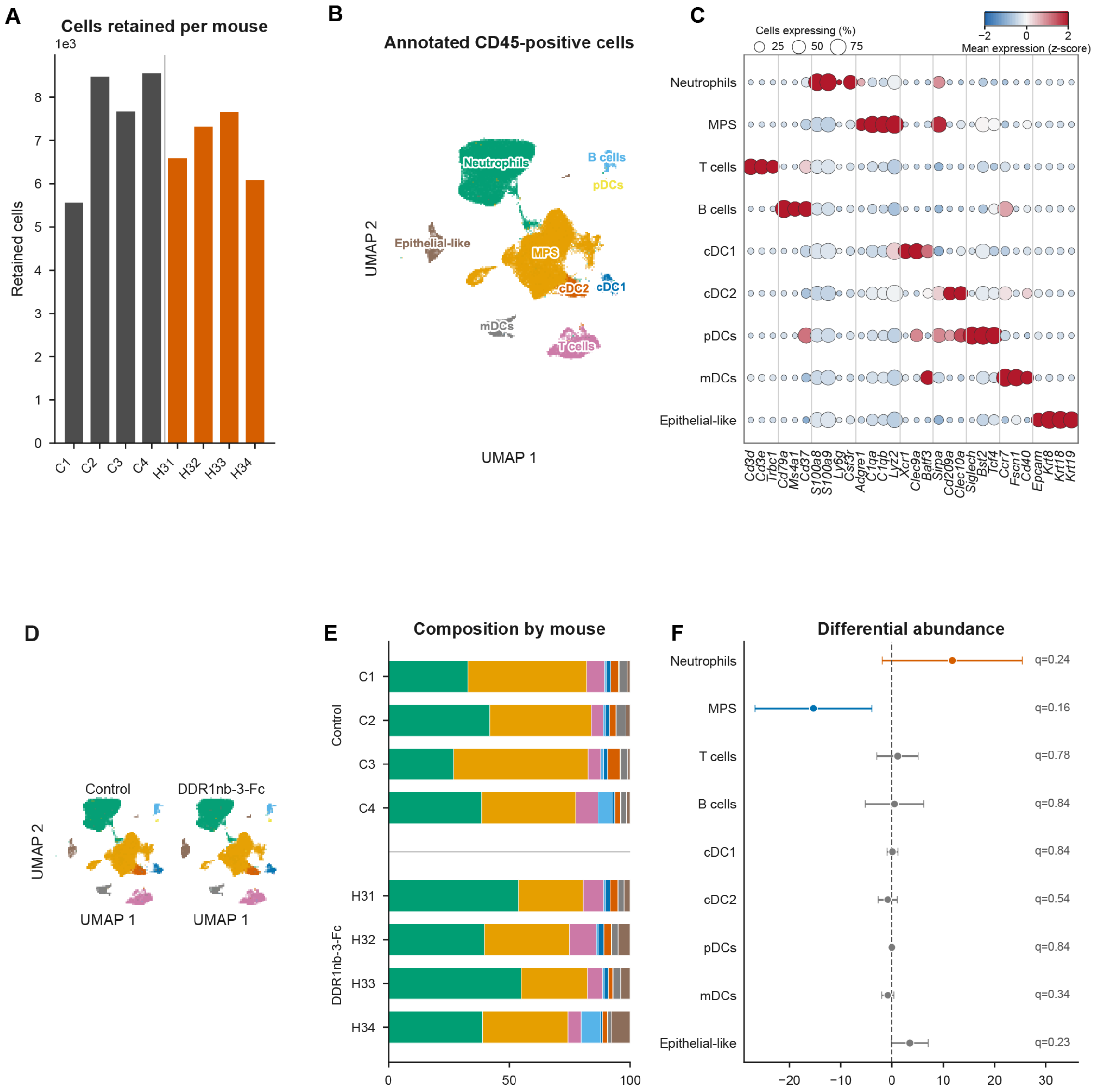


**Supplementary Fig. S15. Annotation and composition of CD45-enriched immune cells from orthotopic KPC1199 tumors.**

(A) Numbers of cells retained from each mouse after preprocessing. Gray bars denote control mice, and orange bars denote DDR1nb-3-Fc-treated mice. (B) UMAP visualization of 57,907 CD45-enriched tumor-associated immune cells assigned to nine major cell classes. (C) Canonical-marker dot plot supporting annotation of the nine major cell classes. Dot size indicates the percentage of cells expressing each gene, and color indicates z-scored mean log-normalized expression across cell classes. (D) UMAP projections shown separately for control and DDR1nb-3-Fc-treated tumors using the same embedding. Equal numbers of cells were sampled from the two treatment groups for visualization only; all retained cells were used for quantitative analyses. (E) Relative abundance of the nine annotated cell classes in each individual mouse. (F) Differences in cell-class frequencies between DDR1nb-3-Fc-treated and control tumors, expressed as treatment-minus-control differences in percentage points. Points indicate mouse-level mean differences, and whiskers indicate 95% confidence intervals. Two-sided Welch's t tests were performed using mice as independent experimental units, with P values adjusted across the nine cell classes using the Benjamini–Hochberg method. Adjusted P values are reported as q values; n = 4 mice per group.

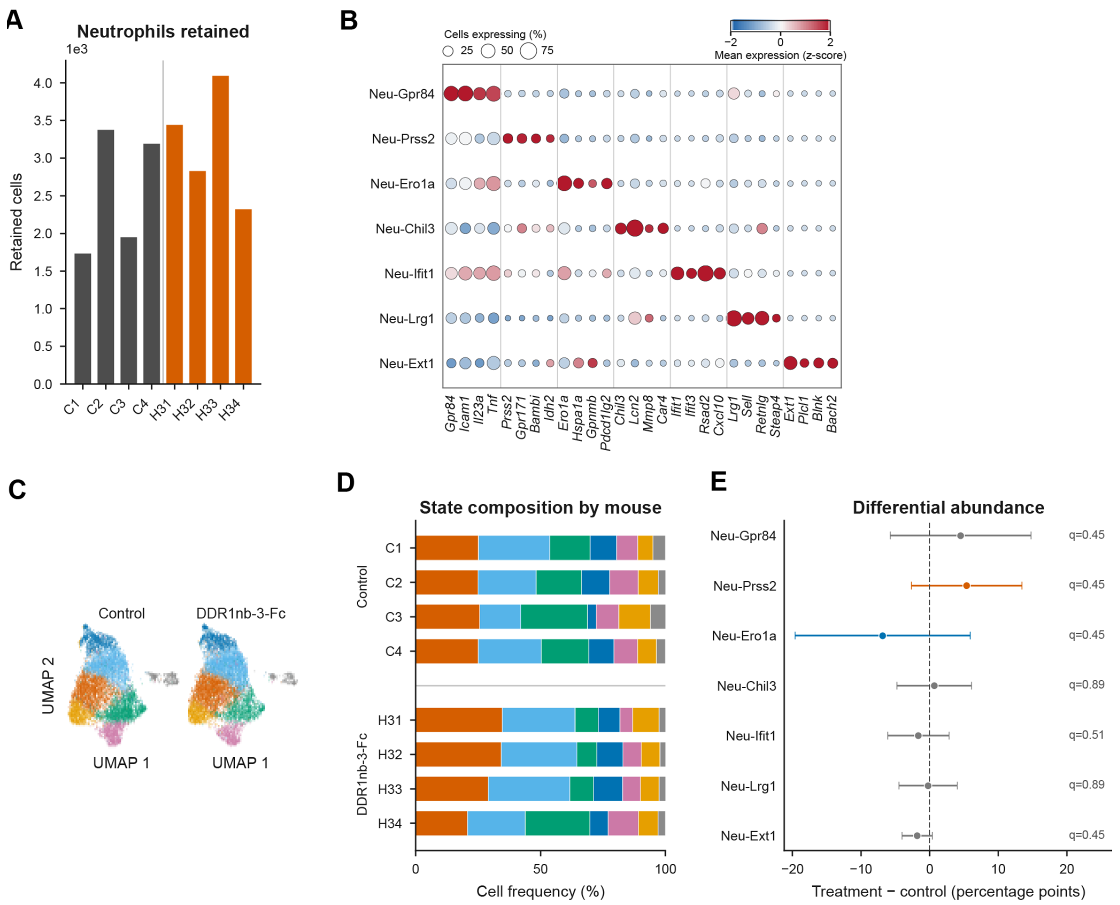


**Supplementary Fig. S16. Marker expression, composition, and between-mouse variation across seven neutrophil states.**

(A) Numbers of neutrophils retained from individual orthotopic KPC1199 tumors. Gray bars denote control mice, and orange bars denote DDR1nb-3-Fc-treated mice. (B) Marker-gene expression across the seven transcriptionally defined neutrophil states. Dot size indicates the percentage of cells expressing each gene, and color indicates z-scored mean log-normalized expression across states. (C) UMAP projections of neutrophils from control and DDR1nb-3-Fc-treated tumors, displayed separately using the same embedding. Equal numbers of cells were sampled from the two treatment groups for visualization only; all retained neutrophils were used for quantitative analyses. (D) Relative contributions of the seven transcriptional states to the neutrophil compartment of each individual mouse. (E) Differences in neutrophil-state frequencies between DDR1nb-3-Fc-treated and control tumors, expressed as treatment-minus-control differences in percentage points. Frequencies were calculated within the neutrophil compartment of each mouse. Points indicate mean differences, and whiskers indicate 95% confidence intervals. Two-sided Welch's t tests were performed using mice as independent experimental units, with P values adjusted across the seven states using the Benjamini–Hochberg method. Adjusted P values are reported as q values; n = 4 mice per group.

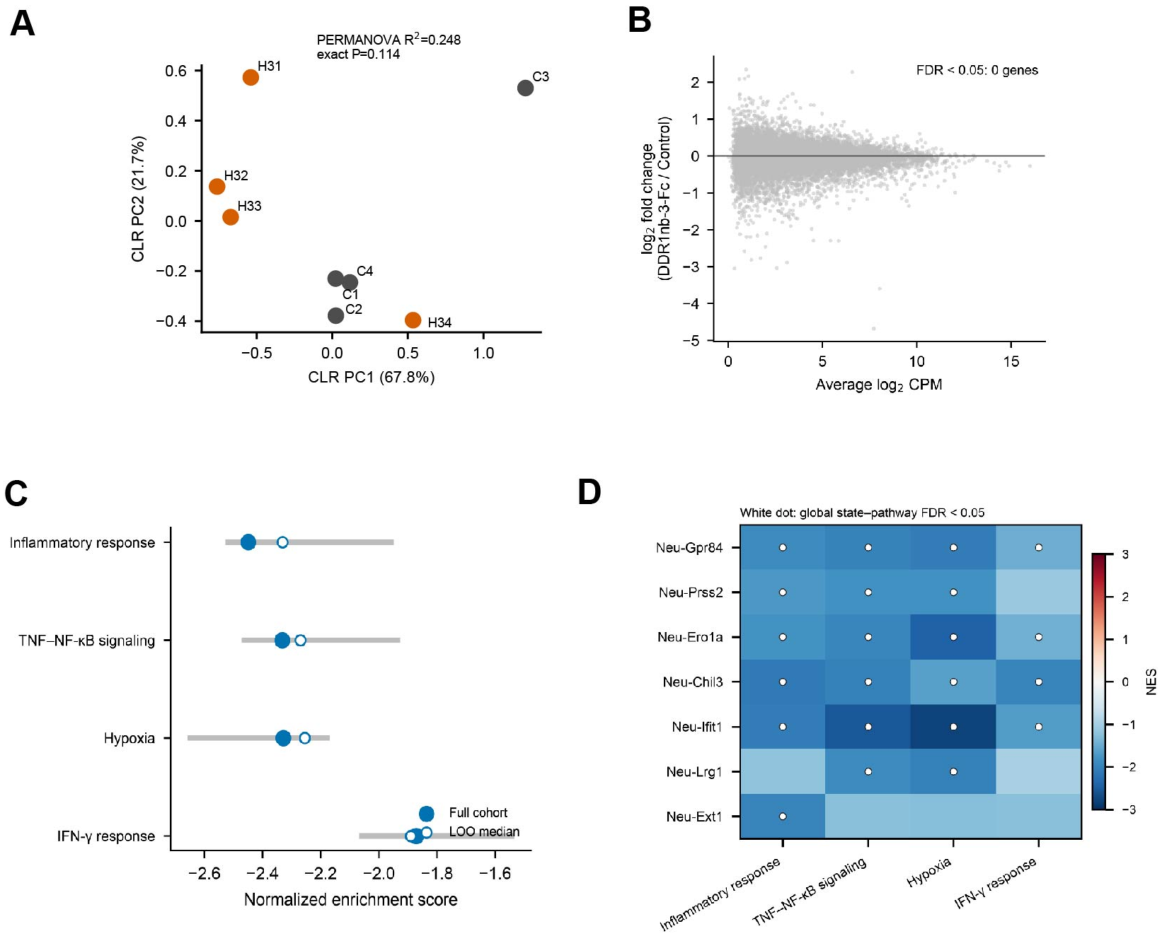


**Supplementary Fig. S17. Mouse-level compositional, pseudobulk, and robustness analyses of tumor-associated neutrophils.**

(A) Principal-component analysis (PCA) of centered log-ratio (CLR)-transformed proportions of the seven neutrophil states across individual mice. The overall treatment-associated difference in neutrophil-state composition was evaluated by exact PERMANOVA using mice as independent experimental units ($R^2 = 0.248$, exact $p = 0.114$). Group dispersion did not differ significantly between treatment groups ($p = 0.971$). (B) MA plot from whole-neutrophil mouse-level pseudobulk differential-expression analysis comparing DDR1nb-3-Fc-treated and control tumors. $Log_2$ fold change is plotted against average $log_2$ counts per million (CPM). No individual gene reached a Benjamini–Hochberg false-discovery rate (FDR) < 0.05. (C) Leave-one-mouse-out sensitivity analysis of the four core treatment-associated pathways identified by whole-neutrophil GSEA. Filled circles indicate NES values obtained from the complete eight-mouse cohort, and open circles indicate the median NES across leave-one-mouse-out analyses. The direction of enrichment was preserved in all eight leave-one-mouse-out analyses for each displayed pathway. (D) State-resolved GSEA showing normalized enrichment scores for inflammatory response, TNF–NF-κB signaling, hypoxia, and IFN-γ response across the seven transcriptionally defined neutrophil states. White dots indicate state–pathway combinations that met FDR < 0.05. The distributed pattern across multiple states does not support attribution of the treatment-associated transcriptional response to a single neutrophil state.

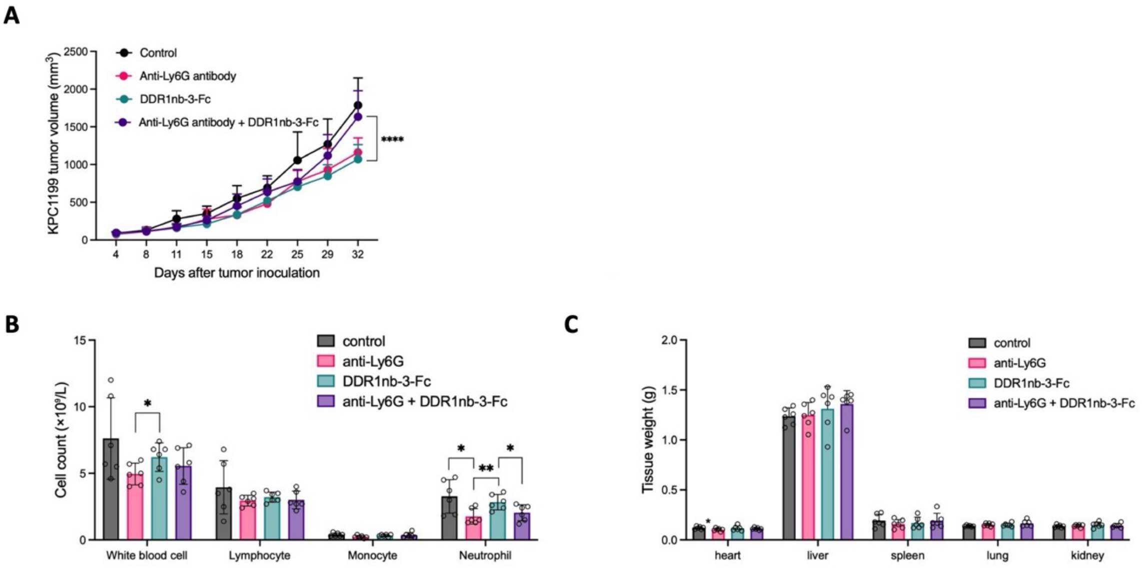


**Supplementary Fig. S18. Longitudinal tumor growth, peripheral blood cell counts, and organ weights in the anti-Ly6G co-treatment experiment.**

(A) Longitudinal tumor-volume measurements from an independent subcutaneous KPC1199 experiment comparing human IgG1 Fc control, anti-Ly6G antibody, DDR1nb-3-Fc, and DDR1nb-3-Fc plus anti-Ly6G antibody; n = 6 mice per group. (B) Peripheral blood white blood cell, lymphocyte, monocyte, and neutrophil counts measured at the study endpoint on day 32 in the same treatment groups. (C) Endpoint wet weights of the heart, liver, spleen, lung, and kidney from the same mice. Points represent individual mice, and summary data are presented as mean ± SD. Statistical comparisons were performed as indicated in the figure. Asterisks denote statistical significance: *$p < 0.05$, **$p < 0.01$, ***$p < 0.001$, ****$p < 0.0001$.

**Supplementary Table S1. BLI-derived binding kinetics of DDR1nb-Fc candidates for human and mouse DDR1-ECD**

| | Human DDR1-ECD | | | Mouse DDR1-ECD | | |
|---|---|---|---|---|---|---|
| | $K_{on}$ ($10^5 M^{-1} S^{-1}$) | $K_{off}$ ($10^{-4} S^{-1}$) | $K_D$ (nM) | $K_{on}$ ($10^5 M^{-1} S^{-1}$) | $K_{off}$ ($10^{-4} S^{-1}$) | $K_D$ (nM) |
| **DDR1nb-3-Fc** | 2.593 | 0.3988 | 0.1538 | 1.718 | 2.354 | 1.370 |
| **DDR1nb-8-Fc** | 2.919 | 13.70 | 4.692 | 3.407 | 7.317 | 2.148 |
| **DDR1nb-13-Fc** | 2.645 | 9.449 | 3.572 | 4.440 | 18.24 | 4.108 |
| **DDR1nb-31-Fc** | 3.020 | 9.443 | 3.127 | 5.280 | 20.68 | 3.916 |
